# Tandem-repeat proteins introduce tuneable properties to engineered biomolecular condensates

**DOI:** 10.1101/2024.04.16.589709

**Authors:** Tin Long Chris Ng, Mateo P. Hoare, M. Julia Maristany, Ellis J. Wilde, Tomas Sneideris, Jan Huertas, Belinda K. Agbetiameh, Mona Furukawa, Jerelle A. Joseph, Tuomas P.J. Knowles, Rosana Collepardo-Guevara, Laura S. Itzhaki, Janet R. Kumita

**Author notes:** these authors contributed equally (Tin Long Chris Ng, Mateo P. Hoare, M. Julia Maristany).

## Abstract

The cell’s ability to rapidly partition biomolecules into biomolecular condensates is linked to a diverse range of cellular functions. Understanding how the structural attributes of biomolecular condensates are linked with their biological roles can be facilitated by the development synthetic condensate systems that can be manipulated in a controllable and predictable way. Here, we design and characterise a tuneable synthetic biomolecular condensate platform fusing modular consensus-designed tetratricopeptide repeat (CTPR) proteins to intrinsically-disordered domains. Trends between the CTPR structural attributes and condensate propensity were recapitulated across different experimental conditions and by *in silico* modelling, demonstrating that the CTPR domain can systematically affect the condensates in a predictable manner. Moreover, we show that incorporating short binding motifs into the CTPR domain results in specific target-protein recruitment into the condensates. Our model system can be rationally designed in a versatile manner to both tune condensate propensity and endow the condensates with new functions.

## Introduction

Biomolecular condensates, also known as membraneless cellular compartments, assemble specific proteins and other biomolecules in a high local concentration through a range of different processes, including phase separation ^1^. The stability and material properties of biomolecular condensates can be modulated by intrinsic parameters, such as the amino acid sequence, post-translational modifications, and the stoichiometry and composition, as well as through the environmental conditions, including heat, pH stress and salt ^2-5^. Biomolecular condensates exhibit a wide range of material properties, comprising low-to-high viscosity liquids, semi-fluid gels, glasses, and solid aggregates. There is growing evidence indicating that the diverse material properties of condensates are significant to their functionality ^6,7^.

Many naturally occurring RNA-binding proteins that form biomolecular condensates combine well-folded globular domains and intrinsically disordered regions (IDRs) with sequences of low complexity (LC). *In vitro* experiments have revealed that LC IDRs are strong positive and negative modulators of the stability of biomolecular condensates, due to their ability to form weak, multivalent interactions ^8-12^. Nonetheless, folded domains, which can establish specific interactions with other biomolecules, are also important regulators of biomolecular condensate properties ^9,11^.

Recent work has focused on understanding how condensate material properties impact biological functions, and several engineered condensate systems have been developed to control the material properties in a tuneable way. These studies have predominantly focused on using IDR regions to drive phase-separation by appending these molecular adhesives to enzymes to characterise their function within the dense phase of a condensate. Such systems have included the use of the hydrophobic protein elastin ^1,13^, short cationic tags ^14,15^ and IDRs derived from naturally occurring phase-separating proteins ^16-18^. Additionally, multivalent protein scaffolds and oligomerisation domains to create engineered condensates have also been explored ^5,7,19,20^.

In this work, we fuse LC domains (LCDs) to folded consensus-designed tetratricopeptide repeat (CTPR) proteins to create a library of chimeric molecules in order to test how changes to the folded domain affect condensate properties. The LCD2 domains were derived from the N- and C-terminal IDR sequences of the Dpb1 DEAD-box ATPase protein, which has been shown to undergo phase separation associated with the formation of processing bodies in yeast ^21,22^. These sequences have been used as molecular adhesives able to drive *in vitro* phase-separation of different globular proteins to create novel microreactors ^17,23^. Condensate formation for these systems can be controlled through protein concentration, pH and salt concentrations, with the LCD2 molecular adhesives showing a broad range of LLPS-inducing conditions ^17^. For folded domains, we used consensus-designed tetratricopeptide repeats (CTPRs) that, unlike globular proteins, adopt an elongated molecular shape comprised of tandem arrays of between 3 and 16 repeats, each composed of a 34 residue helix-loop-helix motif ^24^. Naturally occurring TPRs have the ability to mediate protein–protein interactions via a range of target recognition modes making them useful for a number of processes, including transcriptional regulation, protein folding, cell-cycle control and neurogenesis ^24-26^. Main and co-workers constructed CTPRs through statistical analysis of amino acid propensity at each position of the TPR, and they showed that these artificial proteins were ultra-stable despite the absence of disulphide bonds ^27^. This repetitive modular architecture and high thermodynamic stability make CTPRs a useful tool for protein engineering, where a rational design approach can be used to endow them with additional functionality including specific client recruitment via grafting of single and multiple copies of short linear binding motifs (SLiMs) between adjacent repeats, and arrangement in diverse, precise, and predictable geometries for multivalent and multi-functional display ^28-30^. Using *in vitro* experiments and computational simulations, we demonstrate that engineering different characteristics to the CTPR domains can alter the condensate propensity in a rational way while also enabling specific protein recruitment to the condensates.

## Results

### Design of the LCD2-CTPR system

Based on the molecular adhesive strategy developed by Arosio and co-workers ^17^, we fused LCD2 (N- and C-terminal sequences, Table S1) to the termini of the CTPR domains. Functionalisation of the CTPRs is achieved by grafting SLiMs into one or more of the short (4-residue) inter-repeat loops, whereby even 58 residue-long inserts can be incorporated without causing major disruption of the overall stability of the CTPR protein ^31^. Using this ability to modify CTPR inter-repeat loops, we incorporated a short tetracysteine motif (CCGPCC) between repeats 1 and 2. This motif binds biarsenical dyes like fluorescein arsenical hairpin binder (FlAsH) ^32^. When unbound, the reagent is not fluorescent, but upon interaction with the tetracysteine (TC) motif, a significant increase in fluorescence quantum yield is observed ^33^. The TC-tag/biarsenical dye system is advantageous due to the small size reducing the chance of tag interference with the properties of the protein of interest, and it lends itself to the visualisation of target proteins inside live cells ^15,34-36^. To test whether changes to the CTPR properties can affect the propensity of condensate formation and whether we can endow the condensates with new functions, we created the following CTPR variants: 1) Comprised of either two, three, or four repeats (LCD2-CTPR2, LCD2-CTPR3, LCD2-CTPR4), 2) presenting an altered CTPR surface by changing all lysine residues to arginine residues (LCD2-R-CTPR3), and 3) adding SLiMs that bind specifically to the protein LC3, a protein that is involved in sequestering substrates to the autophagy pathway ^37^. The LC3-interacting regions (LIRs) were inserted between repeats 3 and 4 of LCD2-CTPR4 and allows us to determine whether the condensates can be functionalised to recruit specific target proteins (LCD2-CTPR4-FUNDC1 and LCD2-CTPR4-ATG13) (Fig. 1).

**Figure 1:**
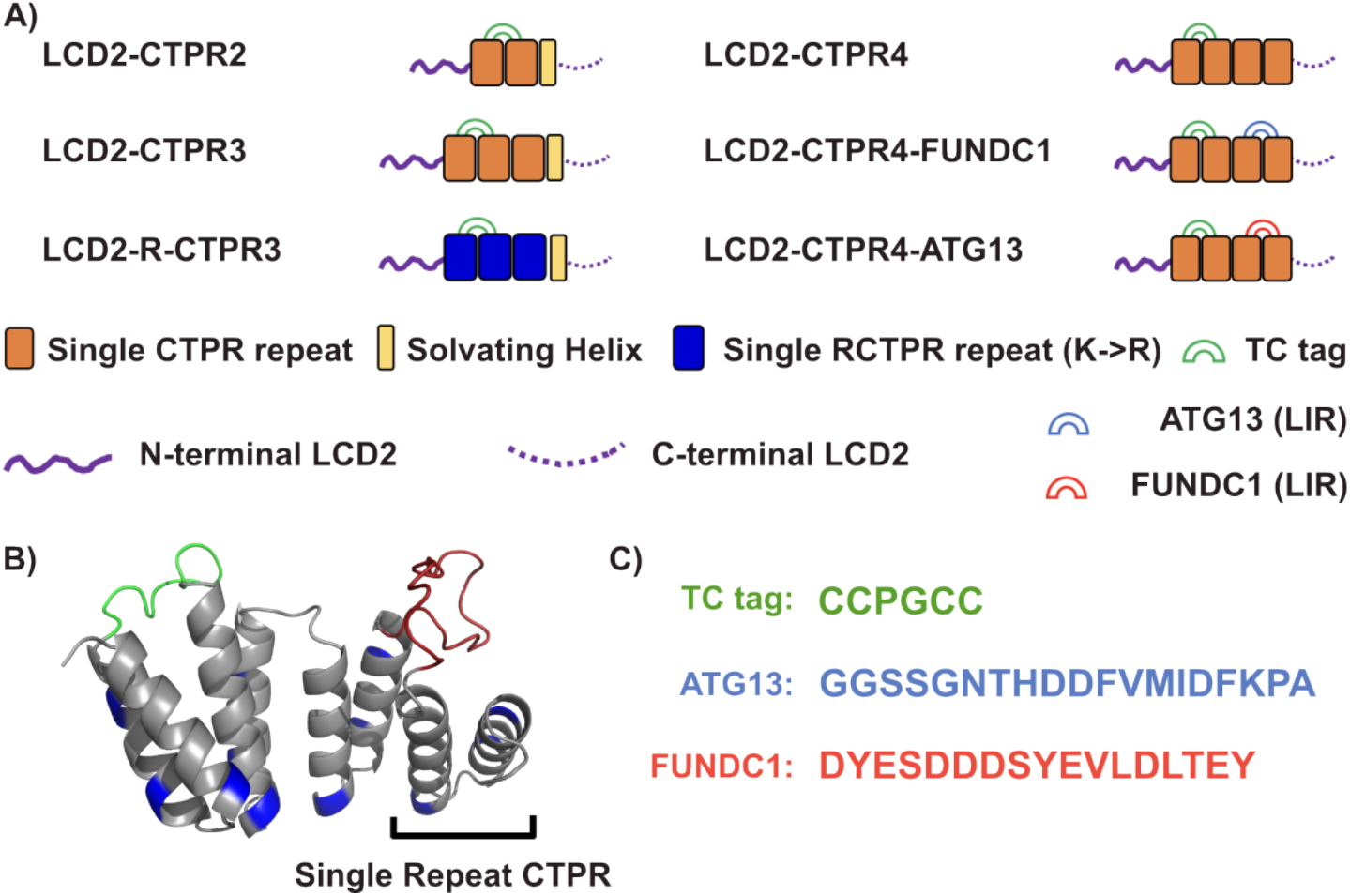
**A)** Composition of different LCD2-CTPR variants. B) AlphaFold 2.0 predicted structure of the 4-repeat CTPR containing a tetracysteine motif (TC, green) between repeats 1 and 2 and an LC3 interacting region (LIR, red) between repeats 3 and 4. The lysine residues, mutated to arginine in the LCD-R-CTPR3 variant, are highlighted in blue. C) Primary sequences of the different endowed loops. Complete primary sequences for all variants are listed in Table S1.

### LCD2-CTPR proteins have high thermodynamic stability and form biomolecular condensates

A four-repeat CTPR protein flanked by N- and C-terminal LCD2 sequences (LCD2-CTPR4) was expressed and purified. The resulting protein could be labelled with FlAsH and, based on mass spectrometry was of high purity (Fig. S1, Table S2). However, unlike the LCD2-GFP protein where high salt concentrations maintained the protein’s solubility ^17^, the LCD2-CTPR4 system phase-separated in 1M NaCl, even with the addition of 1M Urea (Fig. S2*A*). Given that all CTPR variants have high solubility and high stability towards chemical denaturation (Fig. S3, Table S3), with even the smallest, CTPR2 variant having a [GdnHCl]_50_ = 2.58 ± 0.22 M and remaining fully folded in 3M urea (Fig. S4), we introduced 3M urea into the purification buffer to maintain the solubility of the LCD2-CTPR proteins. Using fluorescence microscopy, spherical droplets were observed and formed more readily for LCD2-CTPR4 compared to LCD2-GFP under identical solution conditions (Fig. S2). With this optimised purification method, induced condensate formation using standard dilution methods was achieved ^38^ and used to systematically monitor the condensate-forming properties of the LCD2-CTPR variants. In all LCD2-CTPR variants, we could induce the formation of spherical droplets with different salt concentrations, and these droplets display fusion when imaged in real-time (Fig. 2*A-D*, Fig. S5). Turbidity-based assays ^39^ confirmed the reversibility of the condensates when the samples were heated and cooled (Fig. 2*C*, Fig. S6) or treated with 1,6-hexanediol (Fig. S7). Using this panel of LCD2-CTPR variants, we looked more closely at how the attributes of the CTPR domain can tune condensate propensity and their physico-chemical properties using *in vitro* techniques and *in silico* simulations.

**Figure 2:**
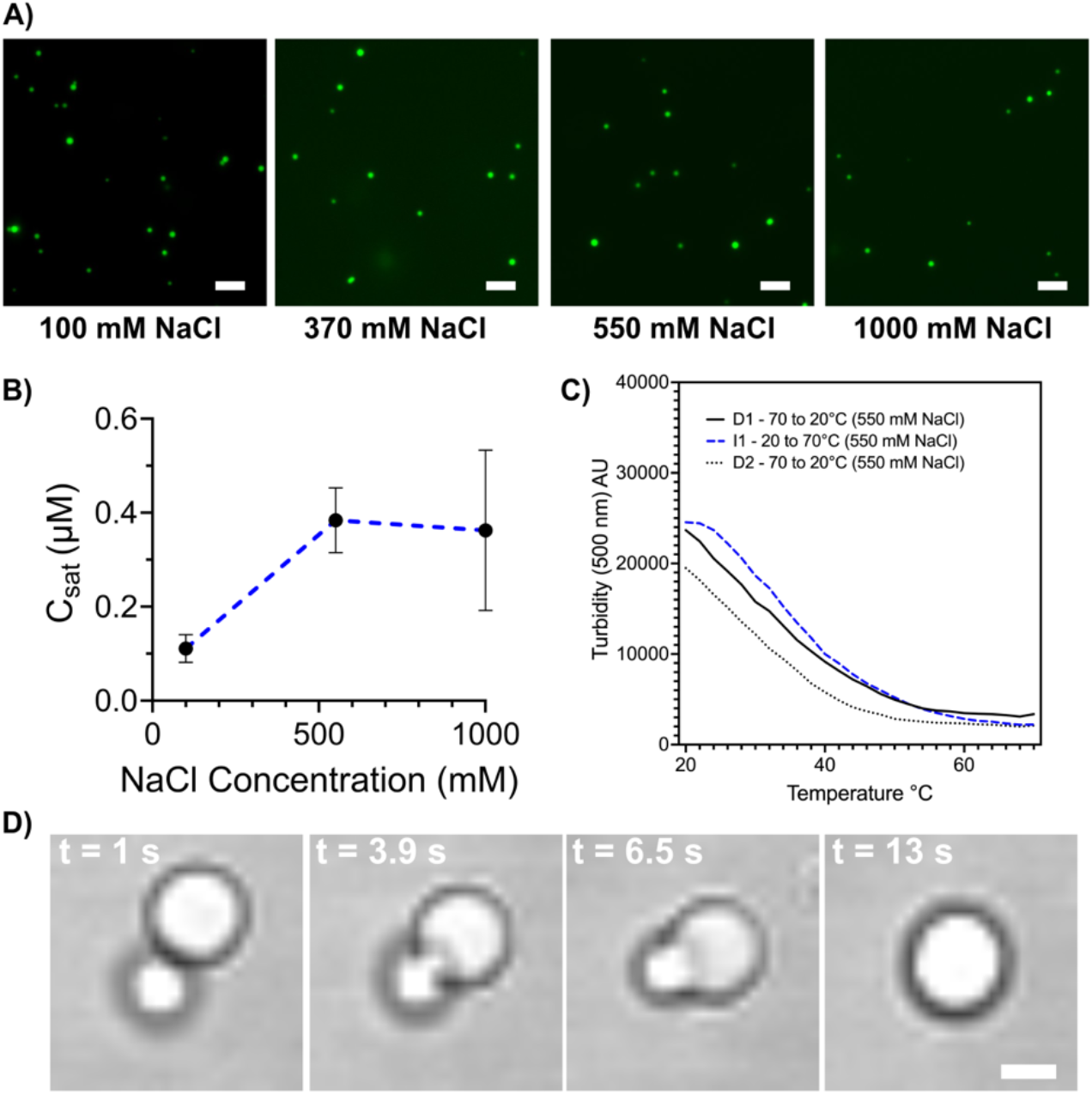
**A)** Fluorescent microscopy images of LCD2-CTPR4 after dilution into 50 mM Tris-HCl pH 7, 300 mM urea, 0.5 mM TCEP and containing varied salt concentrations. Scale = 5 μm. **B)** C_sat_ assay showing the concentration at which LCD2-CTPR4 phase separates in different salt concentration conditions (50 mM Tris-HCl pH 7, 300 mM urea, 0.5 mM TCEP). **C)** Reversible changes in turbidity as condensates, 2 μM with 50 mM Tris-HCl pH 7, 300 mM urea, 550 mM NaCl and 0.5 mM TCEP, were heated and cooled between 20-70ºC. **D)** Real-time microscopy images showing fusion events of LCD2-CTPR4 in 50 mM Tris-HCl pH 7, 300 mM urea, 1000 mM NaCl, 0.5 mM TCEP. Scale = 2 μm.

### CTPR attributes can alter salt-dependent condensate formation

To characterise the tuneability of the LCD2-CTPR condensates, we looked at the effect of salt concentration on their saturation concentrations (*C*_sat_), the concentration at which, under a fixed set of conditions, a protein transitions from the one-phase to the two-phase regime ^16^. Multivalency plays a pivotal role in determining *C*_sat_, as increasing valency notably lowers the *C*_sat_ threshold, initiating phase separation at lower total protein concentrations ^38^. Using a sedimentation-based assay, a range of concentrations of FlAsH-labelled LCD2-CTPR variants were diluted 10-fold in appropriate buffers to induce condensate formation, the samples were centrifuged and the protein in the dilute phase was quantified using SDS-PAGE analysis. A calibration curve of known protein concentrations versus gel-band density was established for this quantification process (Fig. S8). By plotting the total protein concentrations versus the dilute-phase concentrations, a plateau establishes the value of the *C*_sat_ (Fig. S9) and these *C*_sat_ values are shown in Table 1. This was repeated in 50 mM Tris-HCl (pH 7) with different salt concentrations (maintaining a constant urea concentration of 300 mM) to allow the monitoring of salt-dependent changes to the C_sat_ for the LCD2-CTPR variants (Fig. 3, Table 2).

**Table 1:** *C*_sat_ values with standard deviation for the different LCD2-CTPR variants (50 mM Tris-HCl, pH 7, 300 mM urea, 0.5 mM TCEP, room temperature)

| Protein | $C_{\text{sat}}$ ($\mu\text{M}$) | | |
| --- | --- | --- | --- |
|  | 100 mM NaCl | 550 mM NaCl | 1000 mM NaCl |
| LCD2-CTPR2 | $3.48 \pm 0.47$ | $2.01 \pm 0.15$ | $0.96 \pm 0.08$ |
| LCD2-CTPR3 | $2.16 \pm 0.36$ | $1.47 \pm 0.20$ | $1.01 \pm 0.15$ |
| LCD2-R-CTPR3 | $0.48 \pm 0.04$ | $0.17 \pm 0.04$ | $0.27 \pm 0.13$ |
| LCD2-CTPR4 | $0.11 \pm 0.03$ | $0.38 \pm 0.07$ | $0.36 \pm 0.17$ |
| LCD2-CTPR4-FUNDC1 | $0.03 \pm 0.01$ | $0.57 \pm 0.09$ | $2.25 \pm 0.13$ |
| LCD2-CTPR4-ATG13 | $0.09 \pm 0.05$ | $0.52 \pm 0.09$ | $0.51 \pm 0.10$ |

**Table 2:**
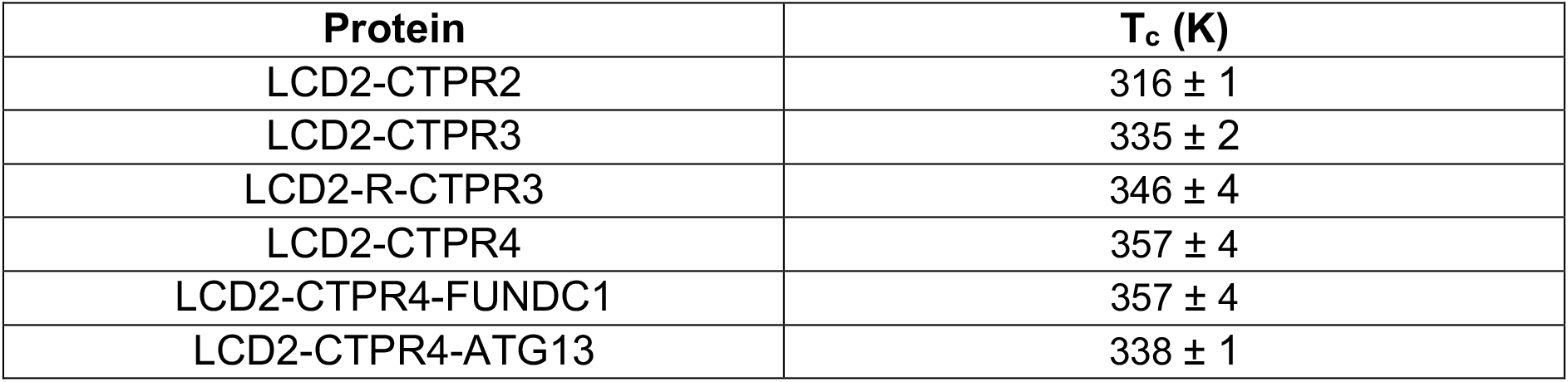
T_c_ values with standard deviation for the different LCD2-CTPR variants (150 mM monovalent salt)

**Figure 3:**
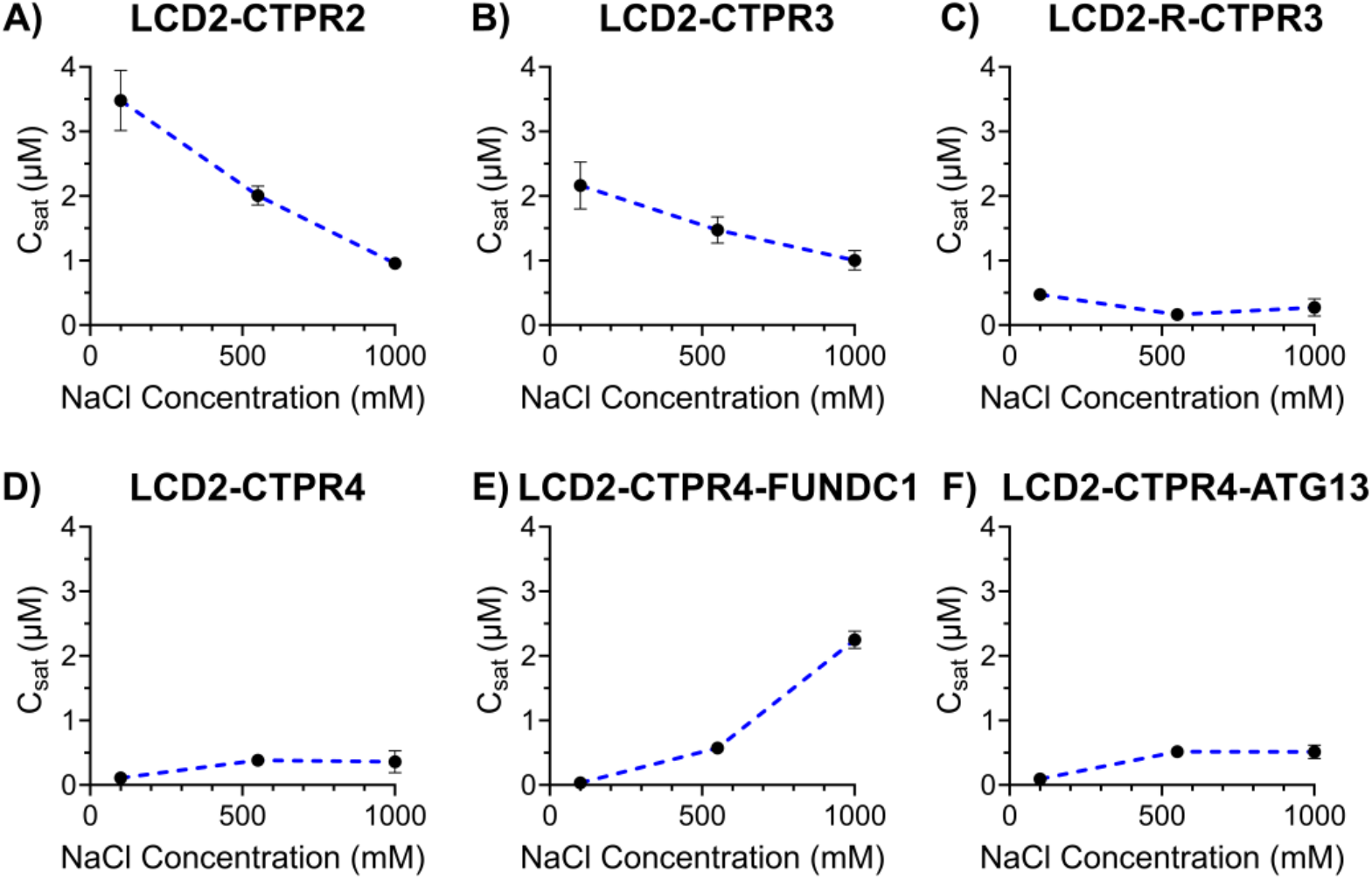
A-F) C_sat_ assay for LCD2-CTPR variants. Plots show how C_Sat_ varies with different salt concentrations in a constant buffer of 50 mM Tris-Cl, pH 7, 300 mM urea, 0.5 mM TCEP. Error is standard deviation (n = 3 to 12).

Comparing the obtained C_sat_ values at the lowest salt concentration (100 mM), we observe a trend in condensate propensity related to the number of CTPR repeat units, where LCD2-CTPR2 is the least prone to form condensates (C_sat_ = 3.48 ± 0.47 μM) followed by LCD2-CTPR3 (C_sat_ = 2.16 ± 0.36 μM), and LCD2-CTPR4 (C_sat_ = 0.11 ± 0.03 μM) being the most condensate-prone. Also, the moderate *C*_sat_ value observed with LCD2-CTPR3 reduce dramatically when lysine-to-arginine substitutions were made in LCD2-R-CTPR3 (C_sat_ = 0.48 ± 0.04 μM). This is consistent with the *in vitro* and *in silico* findings of other research groups, showing that arginine residues can increase phase separation propensity of proteins. This effect is likely due to increased multivalent interactions promoted by the ability of the guanidinium group of the arginine side-chain to establish very energetically favourable cation-π interactions with neighbouring aromatic residues and also strong electrostatic interactions with negatively charged residues ^4,12,40-44^. The incorporation of the LIR-motifs (FUNDC1 and ATG13) did not dramatically alter the C_sat_ values in comparison to the LCD2-CTPR4 (0.11 ± 0.03 μM, 0.03 ± 0.01 μM, 0.09 ± 0.05 μM, respectively). However, when we compare the C_sat_ of these proteins at higher salt concentrations, we observe that increasing salt concentration dramatically inhibits condensate formation of LCD2-CTPR4-FUNDC1, with a C_sat_ of 2.25 ± 0.13 μM in 1M NaCl (Fig. 3*E*, Table 1). This suggests that the negatively charged FUNDC1 loop forms condensate stabilising interactions in low salt concentrations that are masked with high salt concentrations. Furthermore, other salt-dependent trends were observed, such as the clear *C*_sat_ decrease with increasing salt concentration for LCD2-CTPR2 and LCD2-CTPR3 (Fig. 3A, B). These results strongly suggests that the CTPR itself contributes to interactions that are important for modulating condensate propensity.

### Phase diagrams of LCD2-CTPR proteins show that condensate propensity can be systematically modulated

To obtain the phase behaviour across variants under a broader range of solution conditions, we employed PhaseScan, a high-throughput imaging analysis technique. PhaseScan is a combinatorial droplet microfluidic platform designed for rapid, high-resolution, automated analysis of phase separation ^45^. We titrated the FlAsH-labelled protein solution across a range of NaCl and urea concentrations simultaneously (Fig. 4). The dotted line in the phase diagram delineates the phase boundary: above this boundary the protein solution forms a single, dilute phase, and below this boundary the protein solution forms two coexisting phases (dilute and dense). Consistent with our C_sat_ analysis, the LCD2-CTPR variants show clear differences in phase behaviour. Despite the different solution conditions in the PhaseScan, the propensity to form condensates still increases with increasing number of CTPR modules (Fig. 4, *A, B, D*). Second, substituting the surface lysine residues for arginine residues once again shows greatly increased condensate propensity (Fig. 3, *B, C*). The only difference from the C_sat_ trends appears to be for the LCD2-CTPR4 proteins containing two different LC3-binding motifs, where in the PhaseScan analysis introducing these motifs slightly decreases the condensate-forming propensity relative to the unmodified LCD2-CTPR4, with LCD2-CTPR4-FUNDC1 presenting a higher condensate propensity than LCD2-CTPR4-ATG13 (Fig. 3, *E, F*). Overall, it is clear that the CTPR folded domain is a critical regulator of the stability of LCD2-CTPR condensates.

**Figure 4:**
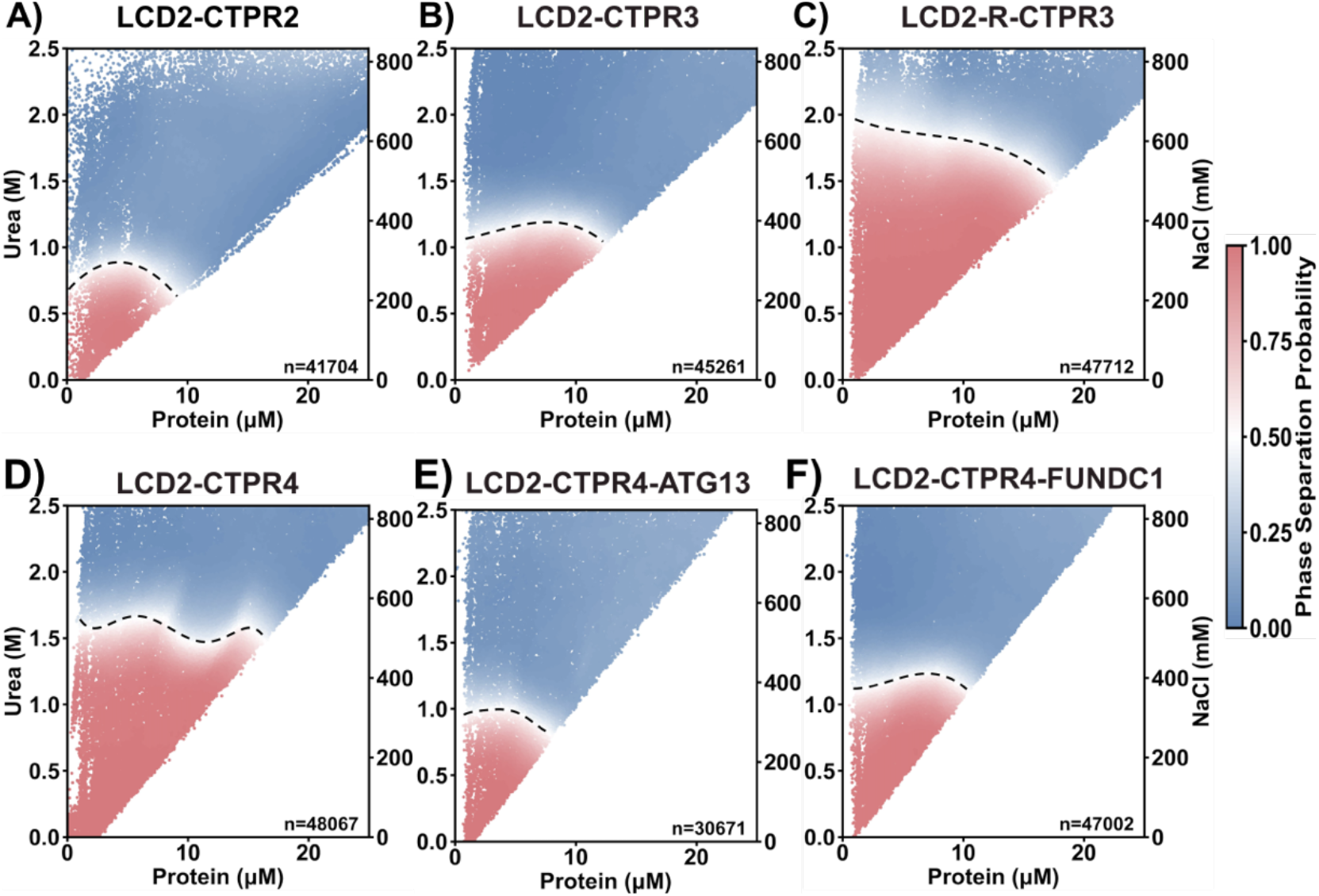
**A-F)** Phase diagrams of LCD2-CTPR variants vs urea and NaCl concentration, 50 mM Tris-HCl pH 8.5, 0.5 mM TCEP. Red and blue data points in the scatter plot correspond to individual microdroplets classified as phase-separated or homogeneous, respectively. The colour-coded heat map shows the estimate phase separation probability over the range of protein vs. urea/salt concentrations. The boundary (dashed line) was acquired by finding a region where the phase separation probability is equal to 0.5 and fitting a line to these coordinates. Data shown is from a single experiment, however, three independent experiments were carried out for each variant to ensure reproducibility.

### *In silico* simulations reveal molecular principles regulating LCD2-CTPR condensate formation

Neither of our experimental systems allow for characterisation of condensate formation in the complete absence of urea, therefore, we use *in silico* simulations of the different LCD2-CTPR variants (sequences in Table S3) to contextualise the *in vitro* analysis. Simulations provide us with a highly controlled environment, in this case the presence of 150 mM monovalent salt (NaCl) that approximates a physiological environment, to enable us to test our hypotheses and explore variables that are difficult to isolate experimentally. These simulations allow us to probe the underlying molecular mechanisms that explain the differential phase separation propensity of the LCD2-CTPR variants.

We used our Mpipi sequence-dependent residue-resolution coarse-grained model for phase-separating proteins, which has been shown to recapitulate experimental temperature-vs-density phase diagrams with near-quantitative accuracy (Fig. 5*A-B*) ^46^. During the simulation, the secondary structure of each separate CTPR module is rigidified. In contrast the IDRs, such as the flanking LCDs and the short loops between CTPR modules, are simulated as fully flexible chains. We perform direct coexistence simulations at various temperatures to compare the stability of the various LCD2-CTPR variants, In a direct coexistence simulation, the condensed diluted phases are simulated in the same elongated box, separated by an interface (Fig. 5*C*)^47-51^. For each temperature probed in the simulations, if after equilibration we detect two phases in coexistence, we measure the density of each phase and plot them versus the simulation temperature to construct coexistence curves (or binodals) in the temperature– density plane (Fig. 5*D*). Our simulations allow us to predict the Upper Critical Solution Temperatures (UCST) of each of our different constructs. That is, the maximum temperature threshold beyond which phase separation is no longer observed is the UCST or critical temperature, *T*_c_, (Fig. 6 *A–F*, Table 2).

**Figure 5:**
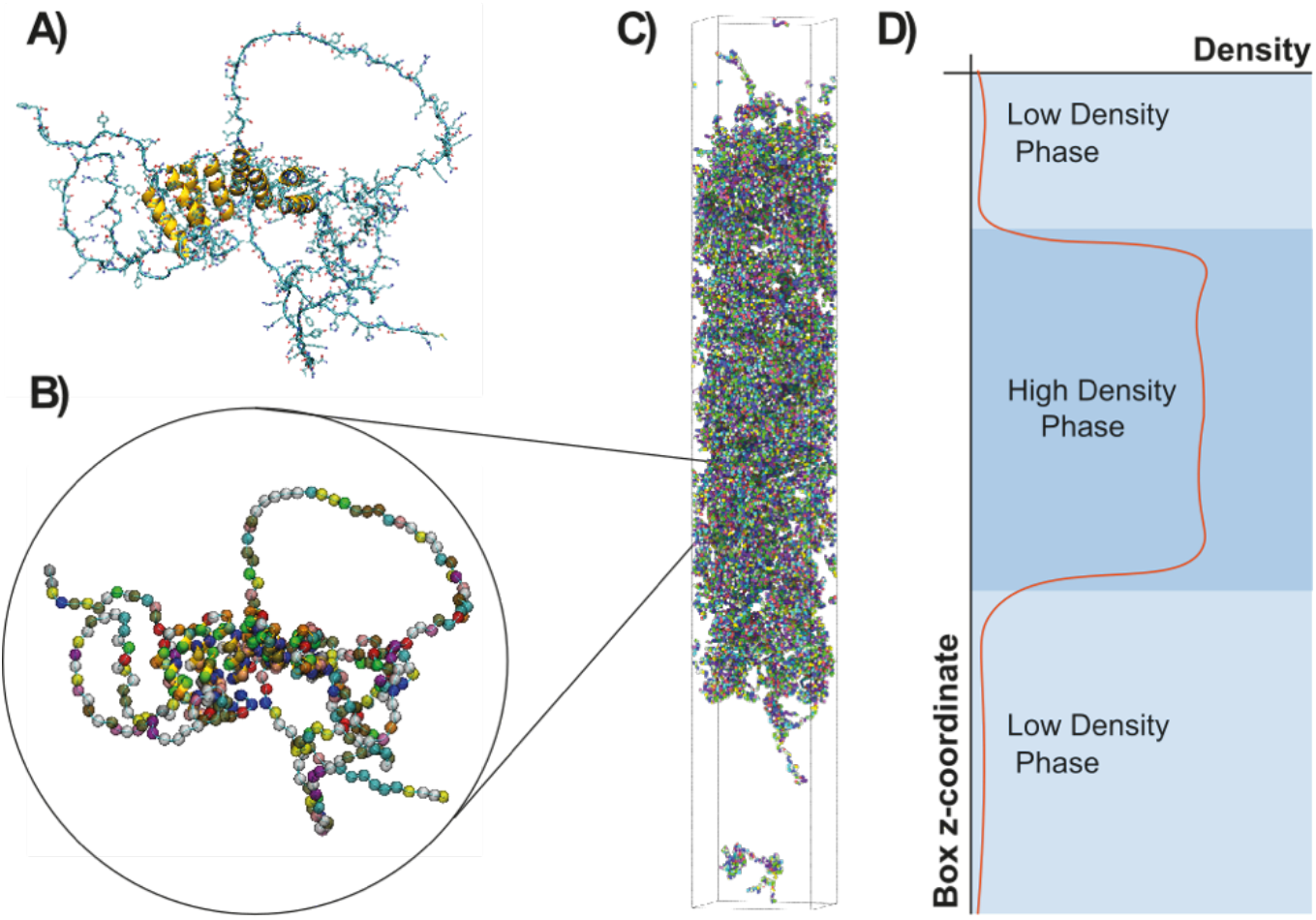
**A)** Predicted structure of a CTPR construct (LCD2-CTPR4) in atomistic resolution, determined using AlphaFold. **B)** Coarse-grained representation of the CTPR-protein. **C)** Simulation box for a direct coexistence simulation. **D)** Profile of the concentration density of the simulated.

**Figure 6:**
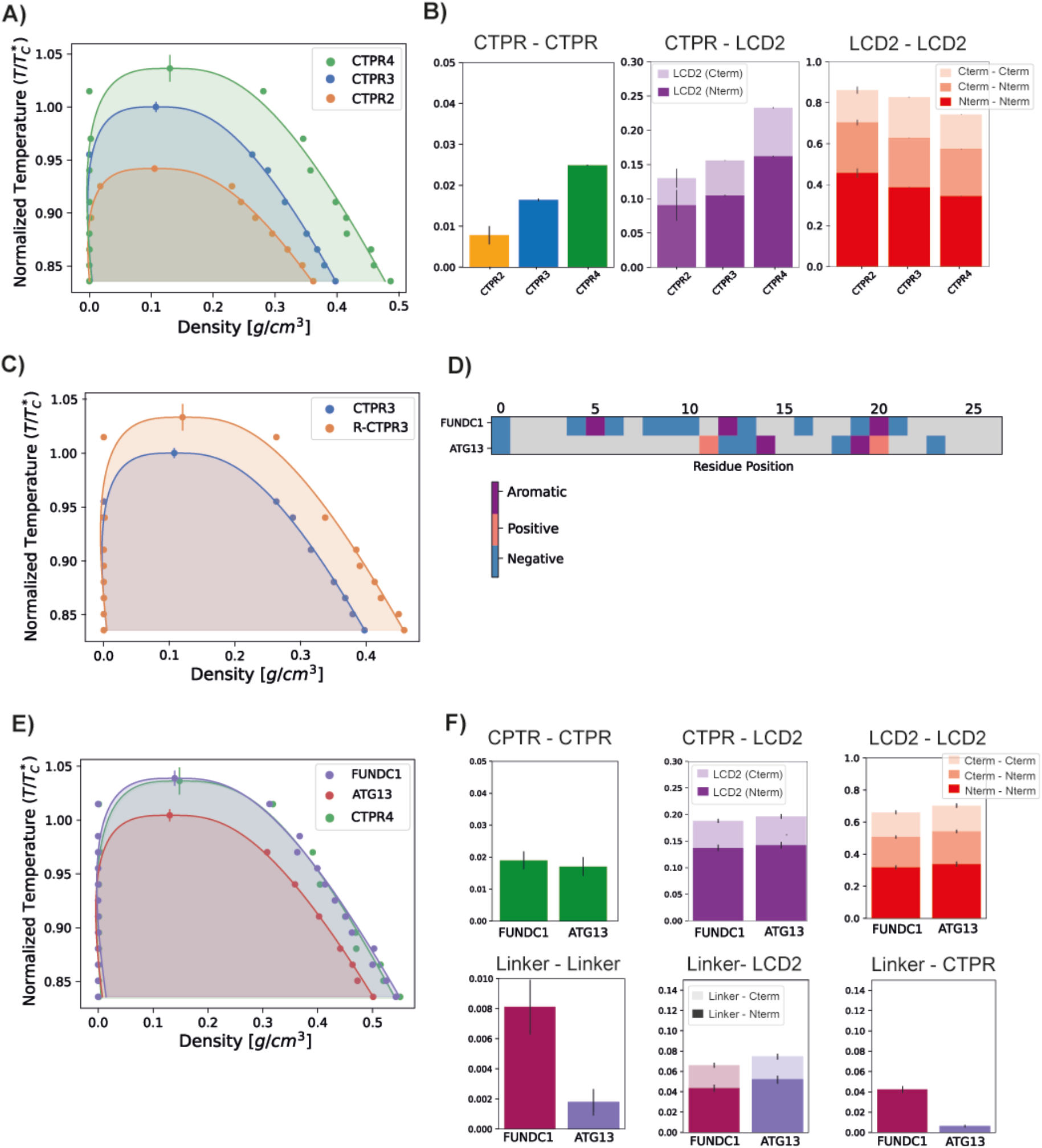
*In silico* analysis of phase-separation of LCD2-CTPR variants. A) Regulation of critical parameters based on the number of CTPR repeats. Binodal in the Temperature-Density phase space of LCD2-CTPR2, LCD2-CTPR3 and LCD2-CTPR4. Temperature in the y-axis is normalized by T*c, the critical temperature of LCD2-CTPR3, to aid in comparison. B) Contact analysis of different protein regions in the condensate: ‘Repeat’ stands for CTPR module, and the LCD2 has been divided into C-termini LCD2 and N-termini LCD2. The contact value represents the percentage of the intermolecular contacts that are of each correspondent type, as an average across simulation frames, and the error bars, the standard error of the average. C) Regulation of critical parameters by selective arginine mutations. Binodal in the Temperature-Density phase space of LCD2-CTPR3 and LCD2-R-CTPR3. D) Analysis of the amino acids in the linkers FUNDC1 and ATG13. E) Regulation of critical parameters according to linker identity. Binodal in the Temperature-Density phase space of LCD2-CTPR4-FUNDC1 and LCD2-CTPR4-ATG13. F) Contact analysis of the different protein regions in the condensate, including the linker region.

In our simulations, LCD2-CTPR2 presented the lowest critical temperature of the set, indicating its general lower propensity for droplet formation (Fig. 6*A*). LCD2-CTPR3 is positioned in the intermediate range. In contrast, the highest critical temperature was observed for LCD2-CTPR4, suggesting the condensates with the strongest thermodynamic stability (Fig. 6*A*). This correlation between increased repeat number and increased critical temperature is in line with the C_sat_ values obtained in 100 mM NaCl (Table 1) and the PhaseScan analysis (Fig. 4*A, B, D*); although, a direct quantitative comparison is not possible due to the differences in solution conditions. In particular for the PhaseScan analysis we varied the concentration of three independent variables – protein, urea and salt, with the latter two changed simultaneously. Our current experimental methods do not allow us to decouple the effects of urea and salt concentration, and a synergistic effect of these components may contribute to the non-linear dependence in the correlation observed experimentally.

One key hypothesis is that increasing the number of globular repeats within the LCD2-CTPR variants enhances the valency of intermolecular interactions ^52-54^. This heightened multivalency is likely to contribute to the formation of a more stable and denser phase. There are many different measures of valency, including computing the total of number residue-residue contacts, or analysing the molecular valency of a given construct ^55^. We focused on the use of residue-residue contacts as this method provides a detailed account of how many intermolecular points of contact exist between proteins, essential for understanding the interaction landscape within the condensate ^56^. An analysis of the most frequent inter-molecular interactions inside the condensate reveals that LCD–LCD interactions contribute most strongly to driving phase-separation of the LCD–CTPR solutions (Fig. 6B). However, we observe that contacts between the LCDs and the CTPRs are clearly observed and that the increased repeat number correlates with increased inter-molecular contacts (Fig. 6B), suggesting that increased valency gives rise to a more densely connected liquid and, hence, enhanced thermodynamic stability of the condensate ^56^. The latter finding confirms that the extended structural fold of the CTPR introduces specific multivalent interactions with the LCDs that can promote phase separation. In addition to the role of multivalency, we hypothesise that the folded domains in the CTPR variants facilitate a more efficient packing density of the proteins within the condensed phase, enhancing its stability. This hypothesis is further supported by the large increase in the critical solution temperature of the LCD2-CTPR3 when the surface lysine residues are substituted for arginine residues to yield the LCD2-R-CTPR3 variant (Fig. 6C).

The differences in condensate critical solution temperature when introducing two different LIR motifs (FUNDC1 and ATG13) between repeats 3 and 4 were also analysed. In accordance with the Phasescan results, LCD2-CTPR4-ATG13 exhibits an decreased propensity for condensate formation when compared to LCD2-CTPR4 and to LCD2-CTPR4-FUNDC1 (Fig. 6E, Table 2). Unlike the experimental results, no observable difference between the critical solution temperature was observed for LCD2-CTPR4 and LCD2-CTPR4-FUNDC1. Despite the linker regions constituting only 5% of the full sequences, contact analysis revealed that the FUNDC1 linker can facilitate intermolecular interactions with the globular domains, both with itself and between them (Fig. 6 F) that were not observed for ATG13. Given that the construct LCD2-CTPR4 has an excess of positive charges, a negatively charged FUNDC1 linker may drive the resulting condensate into electroneutrality and thus, make the dense phase more energetically favourable ^57,58^; this is consistent with our experimental finding that addition of high concentrations of NaCl decreases the C_sat_ of LCD2-CTPR4-FUNDC1 (Fig 3*E*, Table 1).

While our goal with the simulations is to capture and compare the directions of the condensate stability changes upon structural perturbations, it is important to note that the correlation between our *in silico* analysis and the *in vitro* results is qualitative, not quantitative, as our experiments explore the plane of monovalent salt coupled with urea concentrations versus protein concentration, whereas the simulations produce phase diagrams in the plane of temperature. Nevertheless, a robust qualitative agreement is achieved, and the simulations can provide us with deeper mechanistic insights into how specific molecular changes to the CTPR domain, such as mutations of surface arginines or an increasing number of repeats, influence condensate stability. This comparative analysis between *in vitro* phase diagrams and *in silico* condensate propensities confirms that both techniques can be used complementary to predict and influence rational designs into the LCD2-CTPR systems. In both instances, it is clear that the structure and chemical makeup of the CTPR domains highly influences condensate propensity in diverse solution conditions.

### LCD2-CTPR proteins can be readily functionalised for target recruitment into condensates

The CTPRs allow us to easily functionalise the condensates to recruit specific client proteins. Target-binding motifs can be grafted onto the inter-repeat loops, and here, we used LC3 interacting regions (LIRs) that bind to LC3, a member of the ATG8 family of proteins involved in target recognition for the selective autophagy pathway ^37^. Two different LIRs were inserted into the LCD2-CTPR4 protein: the FUNDC1 loop was derived from a protein present in the outer mitochondrial membrane that acts as a receptor for mitophagy in mammals, ^59^ and the ATG13 loop was derived from an autophagy-related protein that regulates phagosome formation ^60^. These LIRs have different LC3-binding affinities, with the LIR from FUNDC1 possessing a higher affinity than that from ATG13 ^61,62^. When grafted into the CTPR4 scaffold without LCD2 regions, the loops do not alter the overall solubility of the proteins and the variants maintain their native fold as monitored by CD spectroscopy (Fig S10A-C), and high native-state stability (Table S2). They also maintain the ability to bind to LC3 as shown in a comparative binding study monitored by western blotting (Fig. S10D).

To study client protein recruitment, condensate formation with the FlAsH-labelled LCD2-CTPR4 variants was induced. Once the condensates were formed, ROX-labelled LC3 protein was added and specific recruitment was monitored by fluorescence microscopy. No co-localisation of the FlAsH-labelled condensates with ROX-labelled LC3 was observed when the LIR-motif is absent (LCD2-CTPR4). The presence of either the FUNDC1 or ATG13 sequences resulted in co-localisation of LC3 within the condensates (Fig. 7*A*). To quantify the extent of this co-localisation, the partition coefficient of ROX-labelled LC3 recruitment to unlabelled LCD-CTPR4 variant condensates was measured (Fig. 7*B*). The data show the FUNDC1 motif resulted in more LC3 co-localisation than the ATG13 motif, which correlates with their relative LC3-binding affinities. This relationship was recapitulated using a sedimentation assay to determine the concentration of LC3 in the dilute phase after co-incubation. It was observed that a lower concentration of LC3 was present in the dilute phase of FUNDC1 compared to the ATG13, suggesting that LCD2-CTPR4-FUNDC1 recruited more LC3 to the dense phase (Fig. 7*C*). These results further demonstrate that not only can we recruit client proteins to our engineered condensates using target-binding loops, but also that the extent of this recruitment can be tuned depending on the binding affinity of the loop used.

**Figure 7:**
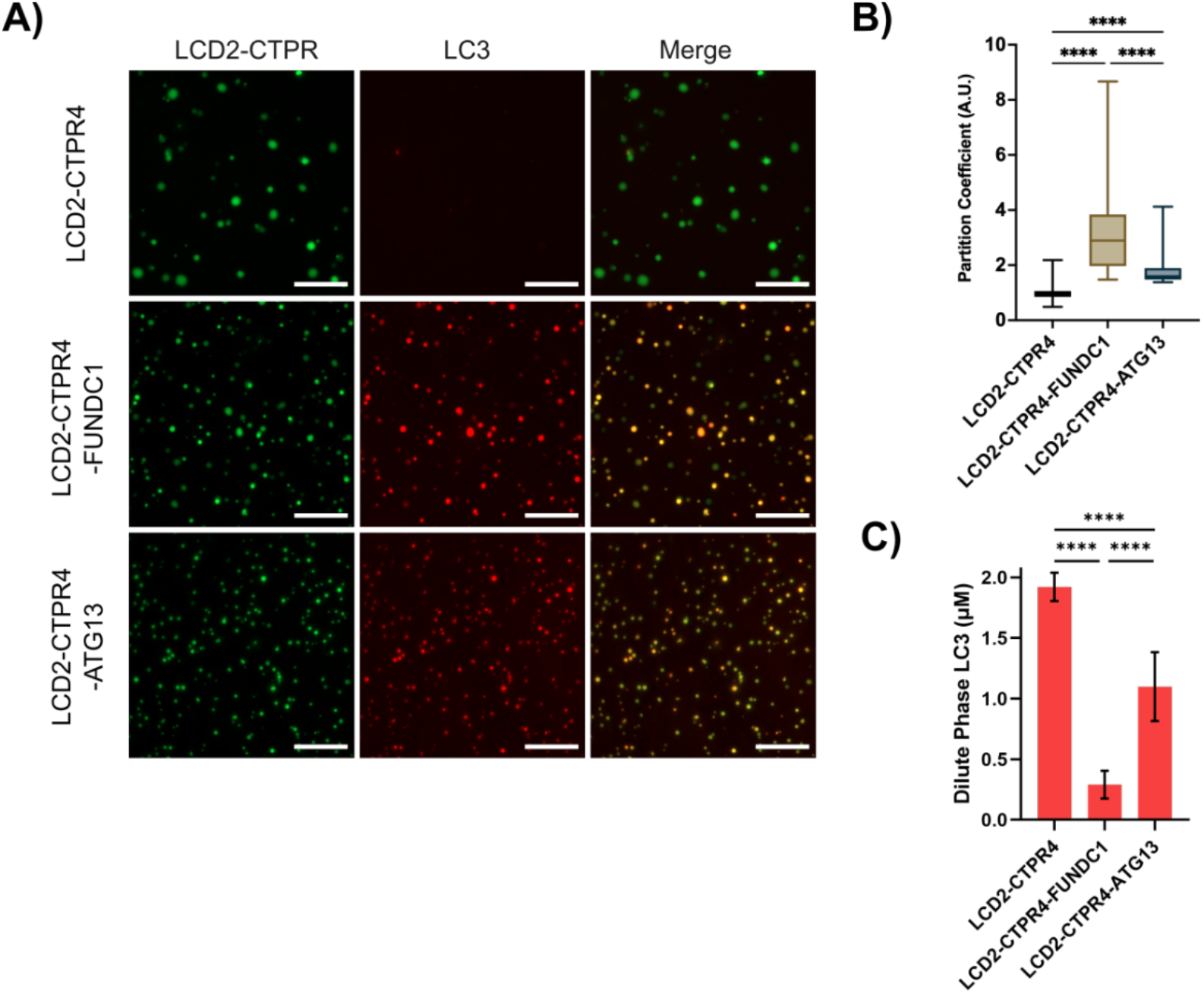
Recruitment of LC3 to LCD2-CTPR4-LIR condensates. A) Fluorescence microscopy with FlAsH-labelled LCD2-CTPR4 variants (50 mM Tris-HCl, pH 7, 100 mM NaCl, 300 mM urea, 0.5 mM TCEP), showing that ROX-labelled LC3 can colocalise to those that contain an LIR. Scale bars are 20 μm. B) Partition coefficient data was measured with unlabelled LCD2-CTPR4 variant condensates (50 mM Tris-HCl, pH 7, 100 mM NaCl, 100 mM urea, 0.5 mM TCEP) co-localised with ROX-labelled LC3. Significance determined by 1-way analysis of variance (ANOVA); **** P < 0.0001. C) Sedimentation assay data showing the concentration of LC3 in the dilute phase of LCD2-CTPR4 samples containing 2 μM total LC3 concentration (50 mM Tris-HCl, pH 7, 100 mM NaCl, 100 mM urea, 0.5 mM TCEP). Error is standard deviation (n = 6), significance determined by 1-way analysis of variance (ANOVA); **** P < 0.0001.

## Discussion

To understand how condensates impact biological functions it is useful to develop synthetic condensates whose properties can be tuned in a controllable and predictable way. Additionally, genetically encodable systems are advantageous so that they can be introduced into the cellular environment. LCD sequences have been used to generate synthetic condensates in bacterial, yeast and mammalian cell systems ^15,18,63-65^ and can be altered to tune the saturation concentrations of phase separation by changes in valency ^18^. Furthermore, they can be rationally redesigned to give rise to more or less viscous liquid condensates or sustain gel-like attributes ^1,19,63^. Systems like PopZ and multivalent *de novo* coiled coils drive phase separation through oligomerisation and can be tuned by altering weak protein-protein interactions, demonstrating applicability in mammalian and bacteria cell systems, respectively ^7,20^. Recently, Arosio and co-workers have described the use of LCD-fusion to promote the phase-separation of globular proteins while maintaining the enzymatic activity of such globular proteins ^17^.

Here we show that fusing LCDs to both termini of a CTPR protein, leads to a phase-separating system where the stability of the LCD2-CTPR condensates is strongly influenced by the unique architecture of the CTPR. Since we have kept the N- and C-terminal LCDs constant in the LCD2-CTPR systems, we can deduce the impact of changes in the CTPR on the phase behaviour of the LCD2-CTPR solutions in a rational manner. As expected, our simulations reveal that the stability of LCD2-CTPR condensates is sustained by numerous associative interactions between the LCDs. Although the CTPR interactions do not drive phase separation (Fig. 6), they introduce LCD to CTPR intactions that allow mutations introduced to this folded region to result in consistent tuneability within the system, irrespective of buffer conditions or condensate induction methods. Using a complementary set of experimental and *in silico* analyses, we have demonstrated that making relatively small changes to the CTPR component can influences condensate formation and physicochemical attributes. Computational analysis indicates that synergistic interactions between CTPR modules and the LCD2 regions are pivotal in promoting phase separation, and with a repeat length of four, there is a strong preference for condensate formation. This prediction is corroborated by our *in vitro* analyses. Interestingly, when we replace lysine residues within the CTPRs modules with arginine residues, we observe a large increase in condensate propensity both *in silico* and *in vitro*. Recent efforts to define the impact of specific amino acids within IDRs that promote or reduce phase separation has demonstrated that saturation of phase separation is inversely proportional to the product of the numbers of arginine and tyrosine residues ^12^. Despite being in the folded CTPR rather than the IDR region, our *in silico* analysis of the residue-to-residue contacts within the LCD2-CTPR variants confirm that the CTPR region has condensate-promoting interactions with the LCDs. Therefore, the lysine-to-arginine mutations introduced likely contribute to increased cation-π and electrostatic interactions that increase overall multivalency and promote phase separation in our system. Lastly, we find that incorporating short functional peptides into the loop regions of the CTPRs alter condensate propensity tuned by their amino acid composition.

The ability to functionalise engineered biomolecular condensates in a facile and versatile manner to enable specific recruitment of target proteins into the dense phase to enhance enzymatic reactions or modulate protein-protein interactions is highly desirable. A variety of strategies to enrich target proteins into engineered condensates have been explored, including short coil-coiled tags and nanobody fusions for GFP or appending directly to a protein of interest to target its partitioning to the condensate phase ^6,7,15,17-20^. A unique feature of the modular CTPR scaffold is that target-binding specificity, valency, and affinity can be readily manipulated, enabling recruitment of one or multiple proteins through engrafting of loops into the inter-repeat loops ^28,29^. As proof-of-concept, we show that incorporating two different LIR-motifs, FUNDC1 and ATG13, results in specific recruitment of LC3 to the dense phase of the condensates. Importantly also, the binding affinity determines the extent of target recruitment.

The relationship between CTPR modulation and condensate-propensity can be rationally understood, on a qualitative level, by comparing *in vitro* experiments and *in silico* modelling. Not only have we demonstrated that there is a tuneable synergy within the LCD2-CTPRs, but the *in silico* analysis of the phase behaviour of these systems closely mirrors our *in vitro* findings. Such a multidisciplinary approach gives us the ability to explore future designs *in silico* in a rational and predictive manner to create novel biomolecular condensates with the ability to specifically recruit target proteins of interest.

## Materials and Method

### LCD2-CTPR plasmid preparation and protein purification

LCD2-CTPR constructs were cloned using gBlock oligos (Integrated DNA technologies) into the pRSET-B vector with FastDigest BamHI and HindIII restriction enzymes (ThermoFisher Scientific (UK) Ltd.) and Anza T4 DNA ligase master mix (Invitrogen). The amino acid sequences of the proteins are listed in Table S3. Recombinant proteins were expressed in chemically competent C41(DE3) *E. coli* cells (Lucigen). Upon reaching OD_600nm_ of 0.6-0.8, bacteria cultures (1L) were induced with isopropyl D-thiogalactopyranoside (IPTG, 0.5 mM, PanReac AppliChem) and incubated (18 h, 20°C, 200 rpm). Cell pellets were collected by centrifugation and resuspended in lysis buffer (50 mM Tris-HCl (pH 8.5), 1 M NaCl, 0.5 mM TCEP, 3 M urea, 1 μg/mL DNase I, 1 cOmplete EDTA-free protease inhibitor tablet (Roche Diagnostics GmbH) for each 50 mL of lysis buffer. Cells were lysed using an EmulsiFlex C5 homogeniser (Avestin) (3-4X, 10000-15000 psi) and centrifuged. The supernatants were purified using HisTrap HP columns (1 mL, Cytiva Ltd.) with Buffer A (50 mM Tris-HCl (pH 8.5), 1 M NaCl, 0.5 mM TCEP with 3 M urea and a linear gradient (8-60% Buffer A + 0.5 M imidazole, 15 column volumes (CV)). LC3 was purified with Buffer B (50 mM sodium phosphate buffer (pH 8.0), 150 mM NaCl (0.5 mM TCEP included for cysteine-variant)) with a linear gradient (8-80% Buffer B + 0.5 M imidazole, 15 CV). LC3 was dialysed (18h, room temperature (RT)) against 50 mM sodium phosphate (pH 8), 150 mM NaCl. LCD2-CTPRs were flash frozen in purification buffer and stored at -80°C. Protein purity was confirmed using SDS-PAGE (Fig. S1) and electrospray ionisation mass spectrometry (Table S1) performed on a Xevo G2 mass spectrometer with data analyzed using MassLynx software (Waters UK) (Yusuf Hamied Department of Chemistry, University of Cambridge, UK).

### FlAsH-EDT_2_ Labelling

LCD2-CTPRs (50 μM) were thawed, centrifuged (2 min, 15000 xg) to remove aggregates and protein was labelled with FlAsH-EDT_2_ (100 μM, Cayman Chemical). Excess dye was removed using a Zeba 7kDa MWCO Spin Desalting Column (Thermofisher) in Buffer A. To determine the concentration of FlAsH-labelled proteins, a correction factor was calculated by measuring the A_280nm_ of the dye divided by the A_max_ of the dye. For FlAsH-EDT_2_ this was determined to be 0.271. Protein concentrations were determined using Eq. 1 where ε_280nm_ was calculated using Expasy ProtParam Tool based on the primary sequence of the protein.

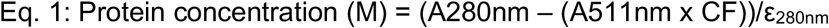

### ROX-maleimide labelling of LC3

LC3 with an N-terminal cysteine was labelled by reacting with a 10x molar excess of TCEP, then adding 2–5x molar excess of ROX-maleimide (Biotium) and incubated (ON, 4°C). 1 mM DTT was added, and excess dye was removed with a 7kDa MWCO Zeba Spin Desalting Column. Protein concentration and labelling efficiency were calculated using Equations 2 and 3:

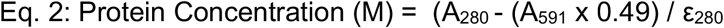

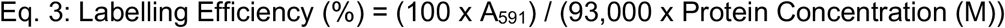

### Fluorescence Microscopy

Imaging of condensate formation was done on a Zeiss AxioVert A1 microscope with a Colibri 7 light source (Carl Zeiss Ltd.), using a LD PN 40x/0.6 Corr objective or an EC PN 63x/1.25 oil objective. Images were captured with a Axiocam 305 mono camera and processed using ImageJ.

### Real-time Microscopy of condensate fusion

Imaging of condensate formation was done on a Leica TCS SP5 DMI6000 B inverted confocal microscope (Cambridge Advanced Imaging Centre, University of Cambridge, UK) with a 100 x 1.4 numerical aperture (NA) HCX PL APO CS total internal reflection fluorescence oil immersion lens with a 512-by-512-pixel photon multiplier tubes (PMTs) as detectors. 2000 frames on each field of view were collected for 30 min.

### PhaseScan

Polydimethylsiloxane (PDMS) devices (Sylgard 184 kit; Dow Corning) were fabricated by casting PDMS on a master wafer made from SU-8 3050 photoresist (A-Gas Electronic Materials Limited) via conventional soft-photolithography methods. Phase diagrams were generated using PhaseScan, a high-throughput phase scanning technology ^45^.

Briefly, stock solutions of different components were mixed at various mass ratios and encapsulated in water-in-oil droplets, individual microenvironments, using a microfluidic device. Fluorescence images of droplets in the observation chamber of the microfluidic device were taken continuously using an epifluorescence microscope (Cairn Research) equipped with 10 × objective (Nikon CFI Plan Fluor 10×, NA 0.3) and high-sensitivity camera (Kinetix, Teledyne Photometrics). Images were analysed via an automated custom Python-based image analysis script ^45^ to detect condensates and to different components within individual microfluidic droplets. The data was plotted as a colour-coded scatter plot, where each point represents individual microfluidic droplets and colour represents the phase separation probability (1 or dark red – phase separated, 0 or dark blue – mixed) assigned to the droplet. This probability was calculated by averaging the phase state assignment or neighbouring droplets. The radius of the neighbourhood was defined as a percentage of data range with an overlayed colour-coded heat map showing the estimated phase separation probability.

### Direct Coexistence MD Simulations

Direct coexistence simulations were performed in a slab geometry, characterised by a simulation box elongated in one of the cartesian coordinates. To coarse-grain the protein system we employed a one-bead-per-amino-acid model, that maps each amino acid of the protein into a single distinct bead. Each amino acid has its particular force field parameters, mass, and charge, to ensure single amino acid resolution. The force field chosen was the Mpipi model, a model optimized for the study of phase-separating systems ^46^. Each CTPR structure was computed through AlphaFold ^66^, optimized by hand via atomistic modelling on Pymol and Gromacs, and finally, systematically cross-validated with published structures and predicted minimized structures. A coarse-graining protocol was then applied, taking the atomistic structure and mapping each bead into the alpha carbons of each protein residue. Globular regions were maintained rigid during the simulation to ensure the conservation of the secondary structure, while intrinsically disordered regions were modeled in a flexible, polymer-like manner.

The simulations were performed using the LAMMPS (Large-scale Atomic/Molecular Massively Parallel Simulator) ^67^. Each phase diagram was computed at several temperature settings, with each set subjected to a thermal equilibrium period of up to 500 ns to guarantee convergence of our results. Each simulation was performed in an NVT ensemble, with a constant number of proteins (N=140) and constant temperature. The specific number of protein copies was carefully chosen to balance computational efficiency and mitigate the impact of finite-size effects. All details of the code are published in ref. (42) and available through github: https://github.com/JuliMaristany/PLD_Scaling_Laws. Due to their size, the trajectories used in this study are fully available upon request in a LAMMPS dump format or an .xyz trajectory file.

### Phase Diagram Fitting

To estimate the critical temperature, we use the law of coexisting densities (Equation 4) ^68^:

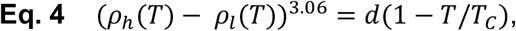

where ρ_*h*_ and ρ_*i*_ are the densities of the protein-rich and protein-depleted phases respectively, 3.06 is the Ising critical exponent, *T*_*C*_ is the critical temperature, and d is a fitting parameter. The law of rectilinear diameter also holds for the estimation of the critical density (Equation 5):

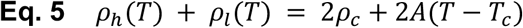

where ρ_*c*_ is the critical density, and A is a fitting parameter.

### Contact Analysis

The number of contacts between the domains of the CTPRs within the slab was analysed using the Visual Molecular Dynamics (VMD) software ^69^. Each protein construct was analyzed firstly by considering each of its distinct domains separately, i.e., the N and C terminal LCD domains, the CTPR repeats, and the designable linkers between CTPRs. For each simulation time frame, we then computed the number of amino acids of a different domain that were at a distance smaller than 7Å from the domain considered. This number, which we refer to as raw contact number, was averaged across frames. Finally, the raw contacts were then normalised by the total number of contacts, giving a resulting percentage. For the specific amino acid sequence of each protein domain, see Table S3. The LCD2 regions were as follows: LCD2-CTPR2: 1-173, 267-334; LCD2-CTPR3 & LCD2-R-CTPR3: 1-173, 301-369; LCD2-CTPR4: 1-173, 320-388; LCD2-CTPR4-ATG13: 1-173, 339-411; LCD2-CTPR4-FUNDC1: 1-173, 337-409. The linker regions were: LCD2-CTPR4-ATG13: 282-308; LCD2-CTPR4-FUNDC1: 282-306.

### Molecular valency

We also conducted contact analysis on the output trajectories from the direct coexistence simulations to determine the frequency of interactions between different CTPR molecules. The analysis was performed on python scripting, coupled to an Ovito API ^70^. The output of the analysis is the number of molecules each CTPR construct is in contact with in each simulation snapshot.

The data pipeline was configured to create virtual bonds with a cutoff of 7 Å, defining all contacts in the trajectory. Those contacts corresponding to different molecules are counted, the output of the analysis is an average of how many different molecules each construct in the protein bulk contacts with. This number is referred to as molecular valency through the text.

### C_sat_ Assays

Proteins were centrifuged to remove aggregates, FlAsH-labelled, buffer exchanged into Buffer A and concentrated. Dilutions to a range of concentrations were made in Buffer A. 10-fold dilutions of these were made in 50 mM Tris-HCl (pH 7.0), varied NaCl, 0.5 mM TCEP buffers, or Buffer A. Final urea concentration in all samples was 300 mM. Samples were centrifuged (5 min, 20,000 xg), and the supernatant was mixed (1:1) with 50 mM Tris-HCl (pH 8.5), 1 M NaCl, 10 M urea, and separated by SDS-PAGE. Band densities were analysed using an Odyssey Fc Imager (LI-COR) and quantified in Image Studio Lite 5.2. Linear regression analysis was carried out on samples diluted 10x in Buffer A (100% dilute phase) to generate a standard curve, which was used to convert the band density of samples to protein concentration. GraphPad Prism 10.0 was employed for plotting and fitting to the model: y = if (mx < c, mx, c), where x is total protein concentration, m an arbitrary constant gradient, and c the C_sat_ value.

### Turbidity Assays

Protein samples were thawed, centrifuged to remove aggregates, buffer exchanged into Buffer A using a Zeba 7kDa MWCO Spin Desalting Column. The samples (20 μM) were diluted 10-fold with 50 mM Tris-HCl (pH 7.0), varied NaCl, 300 mM urea, 0.5 mM TCEP buffer and placed in quartz cuvettes (400 μL) and analysed in a FL 6500 Fluorescence Spectrometer (PerkinElmer LAS (UK) Ltd.). Samples were heated to 70°C then turbidity (excitation and emission set to 500 nm) was recorded as temperature decreased 70°C-to-25°C (D1), then increased 25°C-to-70°C (I2) and a final decrease from 70°C-to-25°C (D2). Readings were taken every 2°C, at 5°C/min, 1.5°C error and 400V PMT. Three independent experimental repeats were performed. For 1,6-hexanediol studies, concentrations of 0%, 4%, or 8% were used. Samples were placed in a 96-well plate (Corning 3881) and turbidity was measured on a FLUOstar Omega Microplate Reader (BMG Labtech). Data were plotted and analysed using GraphPad Prism 10.0.

### LC3 co-localisation Assays

FlAsH-labelled LCD2-CTPRs were diluted to 2 μM in 50 mM Tris-HCl (pH 7.0), 150 mM NaCl, 300 mM urea, 0.5 mM TCEP and ROX-LC3 was added (1:1 ratio). Solutions were imaged using a 63x oil objective and a variable exposure (4–18 ms) for the green channel and 50 ms exposure for the red channel. For partition coefficient assays, unlabelled LCD2-CTPRs (2 μM in 50 mM Tris-HCl (pH 7.0), 150 mM NaCl, 100 mM urea, 0.5 mM TCEP) and ROX-LC3 (1:1 ratio) were combined. Partition coefficients were determined by calculating the fluorescence intensity ratio inside the condensates to that in the surrounding dilute phase. Fluorescence intensities were extracted using ImageJ, applying an intensity threshold to distinguish between the high-intensity condensates and the low-intensity dilute phase. The sedimentation assay was carried out as above, with the addition of ROX-LC3 (1:1 ratio) to the FlAsH-labelled LCD2-CTPRs prior to centrifugation. A control of 2 μM ROX-LC3 in 50 mM Tris-HCl (pH 7.0), 100 mM NaCl, 100 mM urea, 0.5 mM TCEP was used to determine phase proportions for LC3.

## Acknowledgements

The authors would like to thank Dr. Timothy Welsh for his help with initial PhaseScan experiments, Hao Chen, William Murphy and Chloe Burgess for their help with initial condensate experiments, and Dr. Luisa Capalbo, Dr. Lenka Faltova and Prof. Paolo Arosio for insightful discussions. T.L.C.N is supported by a Gates Cambridge International Scholarship, M.J.M. is supported by the Winton Programme for Physics of Sustainability for doctoral funding and B.K.A. is supported by the Cambridge BBSRC-DTP programme. T.S. is supported by the European Union’s Horizon 2020 research and innovation programme under the Marie Skłodowska-Curie grant MicroREvolution (agreement no. 101023060) and the Frances and Augustus Newman Foundation. This project made use of time on HPC granted via the UK High-End Computing Consortium for Biomolecular Simulation, HECBioSim (http://hecbiosim.ac.uk), supported by EPSRC (grant no. EP/R029407/1). J.A.J. acknowledges research support from departmental start-up funds provided by the Department of Chemical and Biological Engineering and the Omenn–Darling Bioengineering Institute at Princeton University. This work was supported by a MRC Career Development Award (MR/W01632X/1, J.R.K.), the Royal Society (RGS\R1\231207, J.R.K.), the Rosetrees Trust (CF2\100013, J.R.K and L.S.I.), the European Research Council (ERC) under the European Union’s Horizon 2020 research and innovation program (grant agreement No 803326 to R.C.G.), the Engineering and Physical Sciences Research Council (EPSRC) (EP/X02332X/1 to J.H.M.), and an ERC grant DiProPhys (agreement ID 10100161, T.P.J.K.).

## Author Contributions

T.L.C.N, M.P.H., M.J.M., R.C-G., L.S.I and J.R.K designed the conceptual framework of the study. T.L.C.N, M.P.H., M.J.M., E.J.W., T.S., J.H., B.K.A., M.F. and J.R.K. performed the experiments. T.L.C.N, M.P.H., M.J.M., E.J.W., T.S., J.H., B.K.A., M.F. J.A.J., T.P.J.K., R.C-G., L.S.I. and J.R.K. contributed to data acquisition and interpretation. T.L.C.N, M.P.H., M.J.M, E.J.W. and J.R.K wrote the manuscript with contributions from all authors.

## References

1 Dai, Y., You, L. & Chilkoti, A. Engineering synthetic biomolecular condensates. Nat Rev Bioeng, 1–15 (2023). 10.1038/s44222-023-00052-6

2 Alberti, S., Gladfelter, A. & Mittag, T. Considerations and Challenges in Studying Liquid-Liquid Phase Separation and Biomolecular Condensates. Cell 176, 419–434 (2019). 10.1016/j.cell.2018.12.035

3 Alberti, S. & Hyman, A. A. Biomolecular condensates at the nexus of cellular stress, protein aggregation disease and ageing. Nat Rev Mol Cell Biol 22, 196–213 (2021). 10.1038/s41580-020-00326-6

4 Krainer, G. et al. Reentrant liquid condensate phase of proteins is stabilized by hydrophobic and non-ionic interactions. Nat Commun 12, 1085 (2021). 10.1038/s41467-021-21181-9

5 Peeples, W. & Rosen, M. K. Mechanistic dissection of increased enzymatic rate in a phase-separated compartment. Nat Chem Biol 17, 693–702 (2021). 10.1038/s41589-021-00801-x

6 Dai, Y. et al. Programmable synthetic biomolecular condensates for cellular control. Nat Chem Biol 19, 518–528 (2023). 10.1038/s41589-022-01252-8

7 Lasker, K. et al. The material properties of a bacterial-derived biomolecular condensate tune biological function in natural and synthetic systems. Nat Commun 13, 5643 (2022). 10.1038/s41467-022-33221-z

8 Feng, Z., Chen, X., Wu, X. & Zhang, M. Formation of biological condensates via phase separation: Characteristics, analytical methods, and physiological implications. J Biol Chem 294, 14823–14835 (2019). 10.1074/jbc.REV119.007895

9 Lin, Y., Currie, S. L. & Rosen, M. K. Intrinsically disordered sequences enable modulation of protein phase separation through distributed tyrosine motifs. J Biol Chem 292, 19110–19120 (2017). 10.1074/jbc.M117.800466

10 Malinovska, L., Kroschwald, S. & Alberti, S. Protein disorder, prion propensities, and self-organizing macromolecular collectives. Biochim Biophys Acta 1834, 918–931 (2013). 10.1016/j.bbapap.2013.01.003

11 Protter, D. S. W. et al. Intrinsically Disordered Regions Can Contribute Promiscuous Interactions to RNP Granule Assembly. Cell Rep 22, 1401–1412 (2018). 10.1016/j.celrep.2018.01.036

12 Wang, J. et al. A Molecular Grammar Governing the Driving Forces for Phase Separation of Prion-like RNA Binding Proteins. Cell 174, 688–699 e616 (2018). 10.1016/j.cell.2018.06.006

13 Quiroz, F. G. & Chilkoti, A. Sequence heuristics to encode phase behaviour in intrinsically disordered protein polymers. Nat Mater 14, 1164–1171 (2015). 10.1038/nmat4418

14 Kapelner, R. A. & Obermeyer, A. C. Ionic polypeptide tags for protein phase separation. Chem Sci 10, 2700–2707 (2019). 10.1039/c8sc04253e

15 Yeong, V., Wang, J. W., Horn, J. M. & Obermeyer, A. C. Intracellular phase separation of globular proteins facilitated by short cationic peptides. Nat Commun 13, 7882 (2022). 10.1038/s41467-022-35529-2

16 Elbaum-Garfinkle, S. et al. The disordered P granule protein LAF-1 drives phase separation into droplets with tunable viscosity and dynamics. Proc Natl Acad Sci U S A 112, 7189–7194 (2015). 10.1073/pnas.1504822112

17 Faltova, L., Kuffner, A. M., Hondele, M., Weis, K. & Arosio, P. Multifunctional Protein Materials and Microreactors using Low Complexity Domains as Molecular Adhesives. ACS Nano 12, 9991–9999 (2018). 10.1021/acsnano.8b04304

18 Schuster, B. S. et al. Controllable protein phase separation and modular recruitment to form responsive membraneless organelles. Nat Commun 9, 2985 (2018). 10.1038/s41467-018-05403-1

19 Garabedian, M. V. et al. Protein Condensate Formation via Controlled Multimerization of Intrinsically Disordered Sequences. Biochemistry 61, 2470–2481 (2022). 10.1021/acs.biochem.2c00250

20 Hilditch, A. T. et al. Assembling membraneless organelles from de novo designed proteins. Nat Chem 16, 89–97 (2024). 10.1038/s41557-023-01321-y

21 Hondele, M. et al. DEAD-box ATPases are global regulators of phase-separated organelles. Nature 573, 144–148 (2019). 10.1038/s41586-019-1502-y

22 Mugler, C. F. et al. ATPase activity of the DEAD-box protein Dhh1 controls processing body formation. Elife 5 (2016). 10.7554/eLife.18746

23 Linsenmeier, M. et al. Dynamics of Synthetic Membraneless Organelles in Microfluidic Droplets. Angew Chem Int Ed Engl 58, 14489–14494 (2019). 10.1002/anie.201907278

24 Main, E. R., Stott, K., Jackson, S. E. & Regan, L. Local and long-range stability in tandemly arrayed tetratricopeptide repeats. Proc Natl Acad Sci U S A 102, 5721–5726 (2005). 10.1073/pnas.0404530102

25 Allan, R. K. & Ratajczak, T. Versatile TPR domains accommodate different modes of target protein recognition and function. Cell Stress Chaperones 16, 353–367 (2011). 10.1007/s12192-010-0248-0

26 Perez-Riba, A., Komives, E., Main, E. R. G. & Itzhaki, L. S. Decoupling a tandem-repeat protein: Impact of multiple loop insertions on a modular scaffold. Sci Rep 9, 15439 (2019). 10.1038/s41598-019-49905-4

27 Main, E. R., Xiong, Y., Cocco, M. J., D’Andrea, L. & Regan, L. Design of stable alpha-helical arrays from an idealized TPR motif. Structure 11, 497–508 (2003). 10.1016/s0969-2126(03)00076-5

28 Diamante, A. et al. Engineering mono- and multi-valent inhibitors on a modular scaffold. Chem Sci 12, 880–895 (2021). 10.1039/d0sc03175e

29 Madden, S. K., Perez-Riba, A. & Itzhaki, L. S. Exploring new strategies for grafting binding peptides onto protein loops using a consensus-designed tetratricopeptide repeat scaffold. Protein Sci 28, 738–745 (2019). 10.1002/pro.3586

30 Perez-Riba, A. & Itzhaki, L. S. The tetratricopeptide-repeat motif is a versatile platform that enables diverse modes of molecular recognition. Curr Opin Struct Biol 54, 43–49 (2019). 10.1016/j.sbi.2018.12.004

31 Ripka, J. F., Perez-Riba, A., Chaturbedy, P. K. & Itzhaki, L. S. Testing the length limit of loop grafting in a helical repeat protein. Curr Res Struct Biol 3, 30–40 (2021). 10.1016/j.crstbi.2020.12.002

32 Martin, B. R., Giepmans, B. N., Adams, S. R. & Tsien, R. Y. Mammalian cell-based optimization of the biarsenical-binding tetracysteine motif for improved fluorescence and affinity. Nat Biotechnol 23, 1308–1314 (2005). 10.1038/nbt1136

33 Pomorski, A. & Krezel, A. Biarsenical fluorescent probes for multifunctional site-specific modification of proteins applicable in life sciences: an overview and future outlook. Metallomics 12, 1179–1207 (2020). 10.1039/d0mt00093k

34 Ng, J. S. W. et al. Using Tetracysteine-Tagged TDP-43 with a Biarsenical Dye To Monitor Real-Time Trafficking in a Cell Model of Amyotrophic Lateral Sclerosis. Biochemistry 58, 4086–4095 (2019). 10.1021/acs.biochem.9b00592

35 Ray, S. et al. alpha-Synuclein aggregation nucleates through liquid-liquid phase separation. Nat Chem 12, 705–716 (2020). 10.1038/s41557-020-0465-9

36 Roberti, M. J., Bertoncini, C. W., Klement, R., Jares-Erijman, E. A. & Jovin, T. M. Fluorescence imaging of amyloid formation in living cells by a functional, tetracysteine-tagged alpha-synuclein. Nat Methods 4, 345–351 (2007). 10.1038/nmeth1026

37 Johansen, T. & Lamark, T. Selective Autophagy: ATG8 Family Proteins, LIR Motifs and Cargo Receptors. J Mol Biol 432, 80–103 (2020). 10.1016/j.jmb.2019.07.016

38 Alberti, S. et al. A User’s Guide for Phase Separation Assays with Purified Proteins. J Mol Biol 430, 4806–4820 (2018). 10.1016/j.jmb.2018.06.038

39 Jung, J. H. et al. A prion-like domain in ELF3 functions as a thermosensor in Arabidopsis. Nature 585, 256–260 (2020). 10.1038/s41586-020-2644-7

40 Bremer, A. et al. Deciphering how naturally occurring sequence features impact the phase behaviours of disordered prion-like domains. Nat Chem 14, 196–207 (2022). 10.1038/s41557-021-00840-w

41 Choi, J. M., Holehouse, A. S. & Pappu, R. V. Physical Principles Underlying the Complex Biology of Intracellular Phase Transitions. Annu Rev Biophys 49, 107–133 (2020). 10.1146/annurev-biophys-121219-081629

42 Maristany, M. J. G. A. A.;Espinosa, J.R.; Huertas, J.; Collepardo-Guevara, R.; Joseph, J.A.. Decoding phase separation of prion-like domains through data-driven scaling laws. bioRxiv (2024). 10.1101/2023.06.14.543914

43 Martin, E. W. et al. Valence and patterning of aromatic residues determine the phase behavior of prion-like domains. Science 367, 694–699 (2020). 10.1126/science.aaw8653

44 Qamar, S. et al. FUS Phase Separation Is Modulated by a Molecular Chaperone and Methylation of Arginine Cation-pi Interactions. Cell 173, 720–734 e715 (2018). 10.1016/j.cell.2018.03.056

45 Arter, W. E. et al. Biomolecular condensate phase diagrams with a combinatorial microdroplet platform. Nat Commun 13, 7845 (2022). 10.1038/s41467-022-35265-7

46 Joseph, J. A. et al. Physics-driven coarse-grained model for biomolecular phase separation with near-quantitative accuracy. Nat Comput Sci 1, 732–743 (2021). 10.1038/s43588-021-00155-3

47 Chapela, G. A. S. G.; Thompson, S.M.; Rowlinson, J.S. Computer simulation of a gas-liquid surface. Part 1. J.Chem.Soc., Faraday Trans. 2 73, 1133–1144 (1977).

48 Espinosa, J. R., Sanz, E., Valeriani, C. & Vega, C. On fluid-solid direct coexistence simulations: the pseudo-hard sphere model. J Chem Phys 139, 144502 (2013). 10.1063/1.4823499

49 Garcia Fernandez, R., Abascal, J. L. & Vega, C. The melting point of ice Ih for common water models calculated from direct coexistence of the solid-liquid interface. J Chem Phys 124, 144506 (2006). 10.1063/1.2183308

50 Ladd, A. J. C. W. L.V. Triple-point coexistence properties of the lennard-jones system. Chem Phys Lett 51, 155–159 (1977).

51 Opitz, A. C. L. Molecular dynamics investigation of a free surface of liquid argon. Phys Lett A 47, 439–440 (1974).

52 Bracha, D., Walls, M. T. & Brangwynne, C. P. Probing and engineering liquid-phase organelles. Nat Biotechnol 37, 1435–1445 (2019). 10.1038/s41587-019-0341-6

53 Li, P. et al. Phase transitions in the assembly of multivalent signalling proteins. Nature 483, 336–340 (2012). 10.1038/nature10879

54 Pappu, R. V., Cohen, S. R., Dar, F., Farag, M. & Kar, M. Phase Transitions of Associative Biomacromolecules. Chem Rev 123, 8945–8987 (2023). 10.1021/acs.chemrev.2c00814

55 Polyansky, A. A., Gallego, L. D., Efremov, R. G., Kohler, A. & Zagrovic, B. Protein compactness and interaction valency define the architecture of a biomolecular condensate across scales. Elife 12 (2023). 10.7554/eLife.80038

56 Espinosa, J. R. et al. Liquid network connectivity regulates the stability and composition of biomolecular condensates with many components. Proc Natl Acad Sci U S A 117, 13238–13247 (2020). 10.1073/pnas.1917569117

57 Chen, S. & Wang, Z. G. Driving force and pathway in polyelectrolyte complex coacervation. Proc Natl Acad Sci U S A 119, e2209975119 (2022). 10.1073/pnas.2209975119

58 Dai, Y. et al. Global control of cellular physiology by biomolecular condensates through modulation of electrochemical equilibria. bioRxiv (2023). 10.1101/2023.10.19.563018

59 Liu, L. et al. Mitochondrial outer-membrane protein FUNDC1 mediates hypoxia-induced mitophagy in mammalian cells. Nat Cell Biol 14, 177–185 (2012). 10.1038/ncb2422

60 Hosokawa, N. et al. Nutrient-dependent mTORC1 association with the ULK1-Atg13-FIP200 complex required for autophagy. Mol Biol Cell 20, 1981–1991 (2009). 10.1091/mbc.e08-12-1248

61 Kuang, Y. et al. Structural basis for the phosphorylation of FUNDC1 LIR as a molecular switch of mitophagy. Autophagy 12, 2363–2373 (2016). 10.1080/15548627.2016.1238552

62 Suzuki, H. et al. Structural basis of the autophagy-related LC3/Atg13 LIR complex: recognition and interaction mechanism. Structure 22, 47–58 (2014). 10.1016/j.str.2013.09.023

63 Garabedian, M. V. et al. Designer membraneless organelles sequester native factors for control of cell behavior. Nat Chem Biol 17, 998–1007 (2021). 10.1038/s41589-021-00840-4

64 Shin, Y. et al. Spatiotemporal Control of Intracellular Phase Transitions Using Light-Activated optoDroplets. Cell 168, 159–171 e114 (2017). 10.1016/j.cell.2016.11.054

65 Wei, S. P. et al. Formation and functionalization of membraneless compartments in Escherichia coli. Nat Chem Biol 16, 1143–1148 (2020). 10.1038/s41589-020-0579-9

66 Jumper, J. et al. Highly accurate protein structure prediction with AlphaFold. Nature 596, 583–589 (2021). 10.1038/s41586-021-03819-2

67 Grindon, C. et al. Large-scale molecular dynamics simulation of DNA: implementation and validation of the AMBER98 force field in LAMMPS. Philos Trans A Math Phys Eng Sci 362, 1373–1386 (2004). 10.1098/rsta.2004.1381

68 Weil, K. G. Molecular Theory of Capillarity. 586 (Clarendon Press, 1984).

69 Humphrey, W., Dalke, A. & Schulten, K. VMD: visual molecular dynamics. J Mol Graph 14, 33–38, 27-38 (1996). 10.1016/0263-7855(96)00018-5

70 Stukowski, A. Visualization and analysis of atomistic simulation data with OVITO-the Open Visualization Tool. Modelling Simul. Mater. Sci. Eng. 18 (2010). 10.1088/0965-0393/18/1/015012

