## Supplementary Information for "Tandem-repeat proteins introduce tuneable properties to engineered biomolecular condensates"

### Supplementary Methods:

#### Circular Dichroism (CD) spectroscopy

CD measurements were conducted with a Chirascan CD spectrometer (Applied Photophysics, Leatherhead, UK) in 1 mm pathlength Precision Cells (110-QS; Hellma Analytics, Müllheim, Germany). Protein samples at 5 µM in sodium phosphate buffer were measured across 205-250 nm wavelengths. Spectra, taken at 1 nm intervals every 0.5 s, were recorded at 20°C, then 90°C, and again at 20°C. Each protein measurement was independently repeated three times, with data baseline corrected and averaged.

#### Gdn-HCl denaturation assay

The different GdnHCl concentrations were prepared by mixing the appropriate volumes of 50 mM sodium phosphate buffer pH 7.5, 150 mM NaCl, 7 M GdnHCl and 50 mM sodium phosphate buffer using a Hamilton Microlab ML510B. The protein concentration used was 15 µM and protein and solutions were dispensed into 96-well, half-area, black polystyrene plates (Corning) and covered with 96-well microplate aluminium sealing tape (Corning). Samples were equilibrated at 25 °C for 2 hr. Emission at 360 nm was measured using a BMG Labtech CLARIOstar® Microplate Reader. The top optic was used in precise mode for 2 cycles with an excitation wavelength of 295 nm and a dichroic PL325 nm filter at 25°C. Each measurement was performed in triplicate and three independent experiments were carried out for each protein variant unless otherwise specified. The data were fit to a two-state model, with denaturation curves fitted directly in GraphPad Prism 10 as described by Perez-Riba et al. (1).

#### LC3 pull-down assay

Ni-Charged MagBeads (GenScript, Oxford, UK) were incubated with 50 µM His-tagged CTPR4 protein (1 hr, RT) then washed with sodium phosphate buffer (x3). The conjugated beads were split into eight aliquots and incubated with purified LC3 at varying concentrations (1 hr, RT). These were washed with TBS-T (50 mM Tris-HCl, 150 mM NaCl, 0.1% (v/v) Tween 20, pH 7.5) (x3) and resuspended in sodium phosphate buffer. The proteins were eluted from beads with phosphate buffer with 500 mM Imidazole. Samples were run on 15% polyacrylamide gels for SDS-PAGE. The proteins were transferred onto a Immobilon-P polyvinylidene difluoride (PVDF) membrane (Merck Millipore) using a Pierce Power Blot Cassette (25 V, 1.3 A, 10 min). The membrane was blocked (5% Dried Skimmed Milk (Marvel) TBS-T, 1 hr), then incubated with a rabbit anti-LC3B primary antibody (GeneTex GTX127375, 1:1000 in 5% milk/TBS-T, ON, 4°C). The membrane was washed with TBS-T (3x 5 min) and incubated with swine anti-rabbit IgG / HRP secondary antibody (Dako P0399, 1:5000 in 5% skimmed milk/TBS-T) (1 hr). The membrane was washed with TBS-T (3x 5 min) and detected using Amersham ECL™ Western Blotting Detection Reagents (Cytiva), imaged using an Odyssey Fc Imager (LI-COR, Lincoln, NE, USA).

### References:


1. A. Perez-Riba, L. S. Itzhaki, A method for rapid high-throughput biophysical analysis of proteins. *Sci Rep* **7**, 9071 (2017).

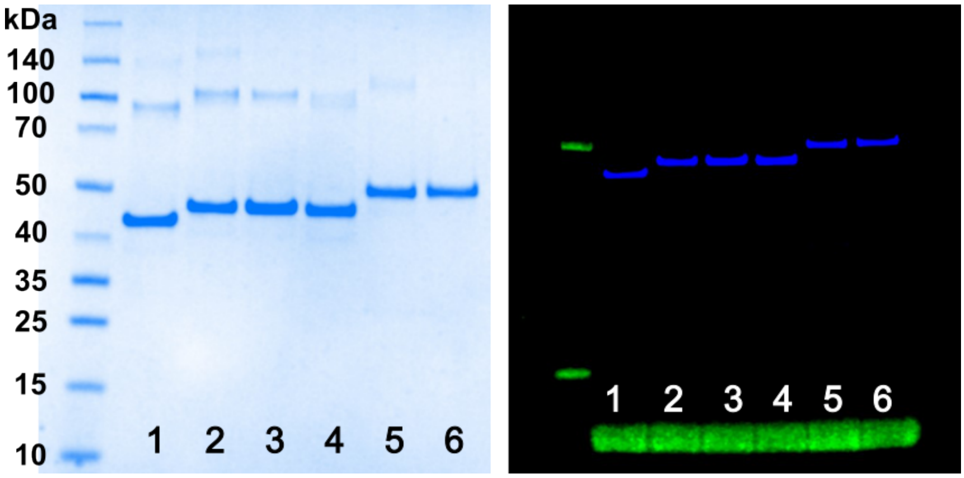


**Fig. S1. SDS-PAGE analysis of purified LCD2-CTPR variants**

A) Images of SDS-PAGE gels of purified LCD2-CTPR proteins. Left: Coomassie stained, Right: FlAsH labelled, 1: LCD2-CTPR2, 2: LCD2-CTPR3, 3: LCD2-R-CTPR3, 4: LCD2-CTPR4, 5: LCD2-CTPR4-ATG13, 6: LCD2-CTPR4-FUNDC1.

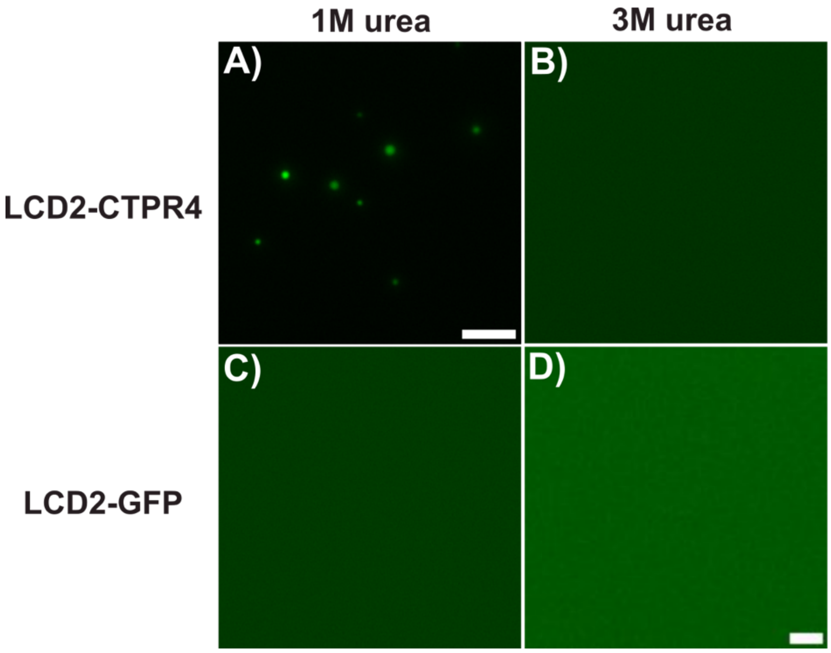


**Fig. S2. Phase separation comparison between LCD2-CTPR4 and LCD2-GFP variants in different buffer conditions.** Microscope images showing phase separation at 2 μM of (A, B) LCD1-CTPR4 and (C, D) LCD2-GFP in two different buffer conditions. (A, C) 50 mM Tris-HCl, pH 8.5, 1 M NaCl, 1 M urea, 0.5 mM TCEP. (B, D) 50 mM Tris-HCl, pH 8.5, 1 M NaCl, 3 M urea, 0.5 mM TCEP. Scale bar = 5 μm.

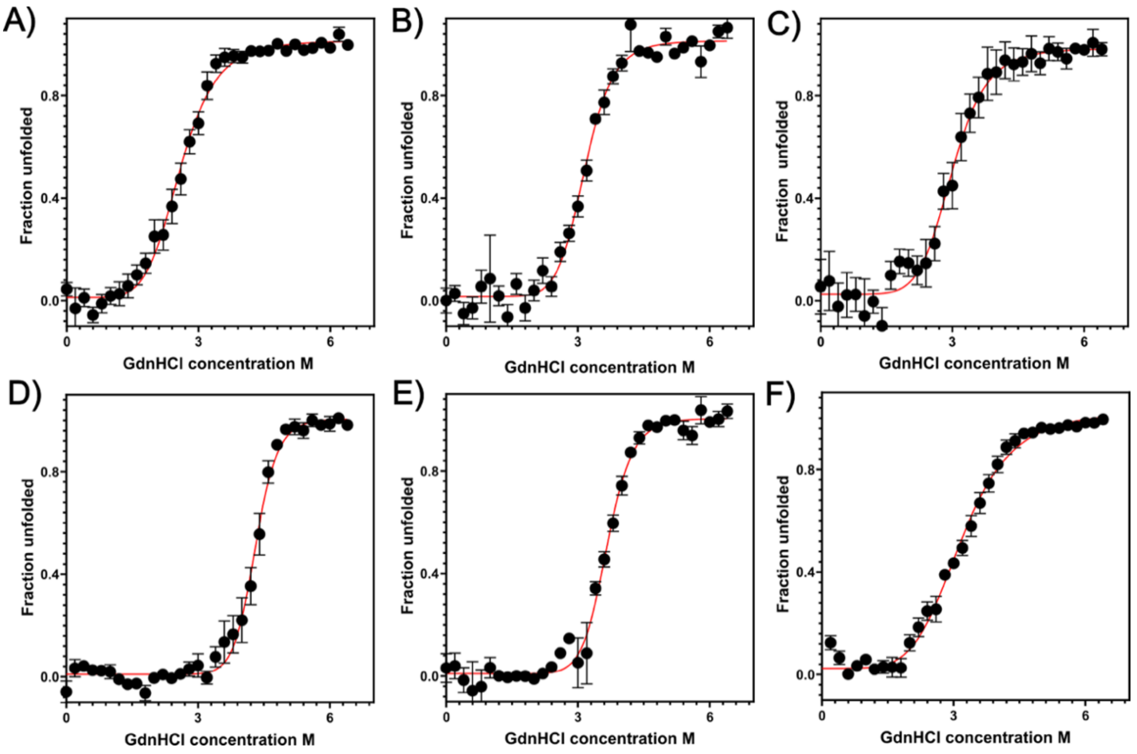


**Fig. S3. GdnHCl-induced denaturation of CTPR-TC proteins.** Purification of CTPR-TC variants yielded samples with concentrations between 150-400 μM. A) CTPR2, B) CTPR3, C) R-CTPR3, D) CTPR4, E) CTPR4-FUNDC1 and F) CTPR4-ATG13. 1 μm of protein in 50 mM sodium phosphate, 150 mM NaCl, pH 7.5 with ranging GdnHCl concentration (from 0 M to 6.4 M) was excited at 295 nm at 25°C. Emission intensity was measured at 360 nm. A minimum of three independent experiments were recorded for each protein. The data were fitted with a two-state model to give the midpoint of unfolding and *m* value (a constant proportional to the change in solvent-accessible surface area upon unfolding). Error bars show the standard error of the mean.

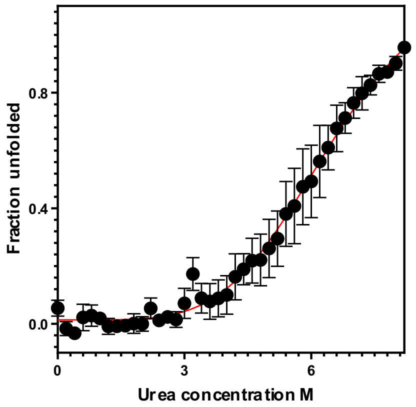


**Fig. S4. Urea-induced denaturation of CTPR2.** Urea-induced denaturation of CTPR2. 1 μm of protein in 50 mM sodium phosphate, 150 mM NaCl, pH 7.5 with urea concentration from 0 M to 8 M was excited at 295 nm at 25°C. Emission intensity was measured at 360 nm. Two independent experiments were recorded for this protein. The data were fitted with a two-state model to give the midpoint of unfolding and *m* value. Error bars show the standard error of the mean.

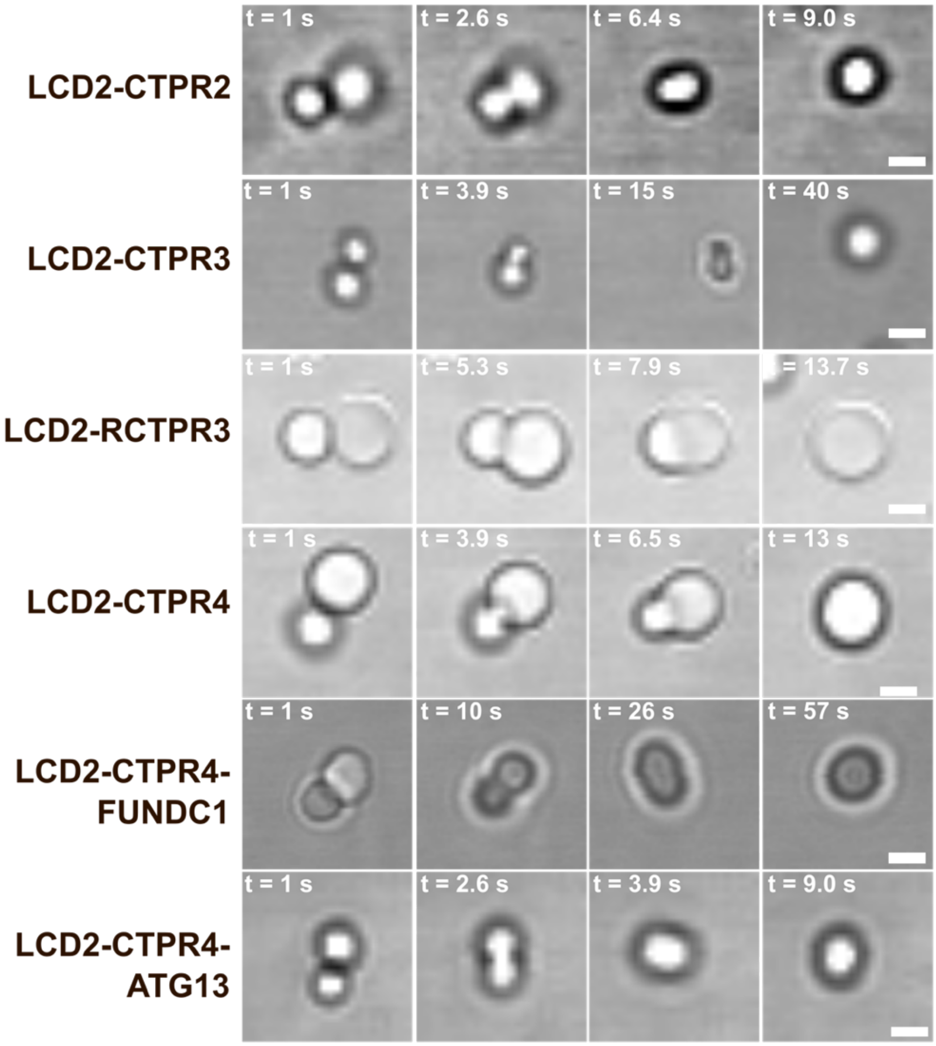


**Fig. S5. Coalescence of the LCD2-CTPR variant condensates.** 2 μM of each LCD2-CTPR variant was monitored in real-time in 50 mM Tris-HCl buffer at pH 8.5, 1000 mM NaCl. Scale bar is 2 μm.

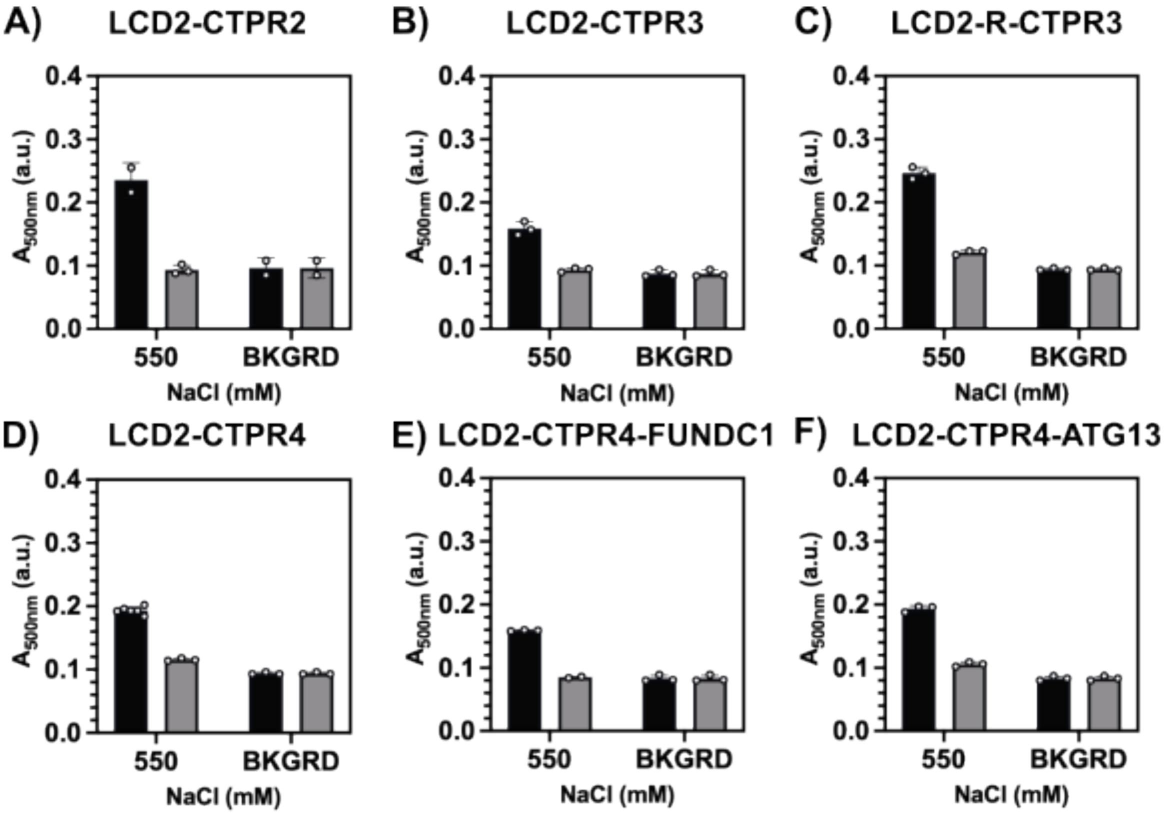


**Fig. S6. 1,6-Hexanediol dissolution of LCD2-CTPRs constructs.** Bar graphs (A-F) show the $A_{500nm}$ for 2 μM LCD2-CTPR variants in 50 mM Tris-HCl, pH 7.0, with 100 mM or 550 mM NaCl, with (grey bars) and without (black bars) 4% 1,6-hexanediol (16H) treatment. Background absorbance is also shown as BKGRD. Significance determined by 2-way analysis of variance (ANOVA); **** $P < 0.0001$, *** $P < 0.0002$, ** $P < 0.0021$, * $P < 0.0332$, ns < 0.1234.

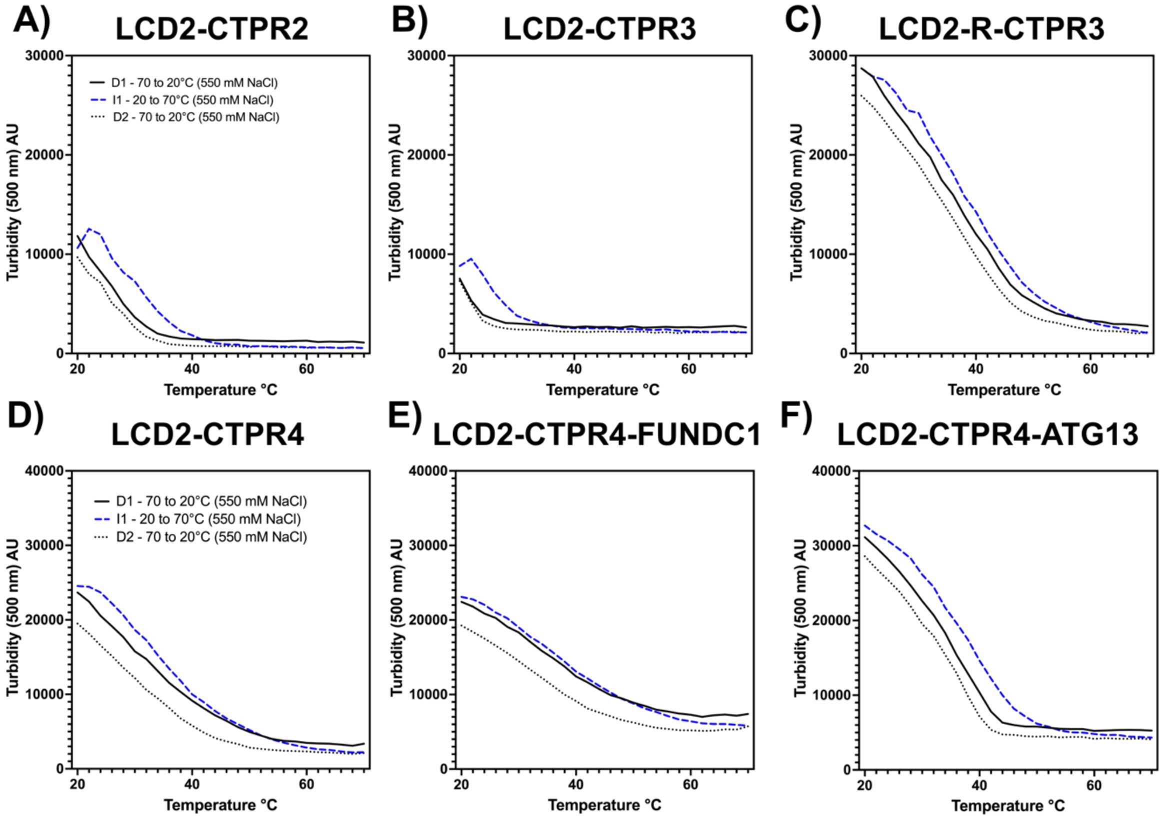


**Fig. S7. Turbidity assay for LCD2-CTPR variants.** Condensates were induced using standard dilution methods and heated to 70°C. The turbidity (at 500 nm) was monitored incrementally from 70 → 20°C (D1: solid black line), followed by an incremental increase from 20 → 70 °C (I1: dashed blue line) and finally another incremental decrease from 70 → 20°C (D2: dotted black line). Samples contained 2 μM protein, 50 mM Tris-HCl pH 7, 300 mM urea and 500 mM NaCl. Data is representative traces from n=3 independent experimental repeats.

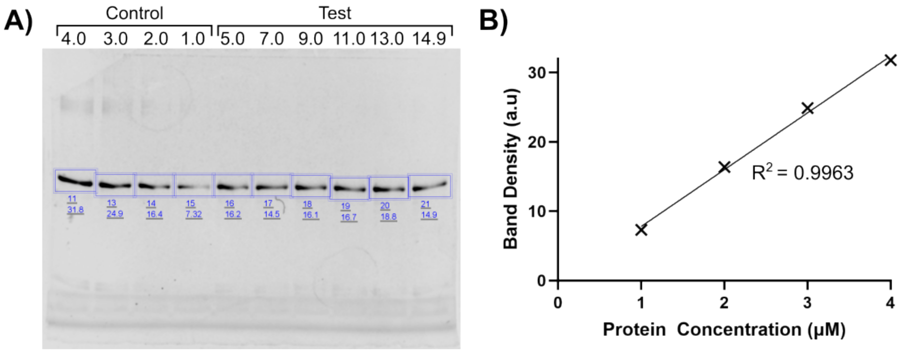


**Fig. S8. Representative $C_{Sat}$ Assay for LCD2-CTPR2 in 50 mM Tris (pH 7), 550 mM NaCl, 300 mM Urea, 0.5 mM TCEP.** A) SDS-PAGE with band density quantification and total protein concentrations (μM) in control and test buffers. B) Linear regression of band density vs protein concentration (determined by UV-vis absorbance) in the control samples, R2 = 0.9963.

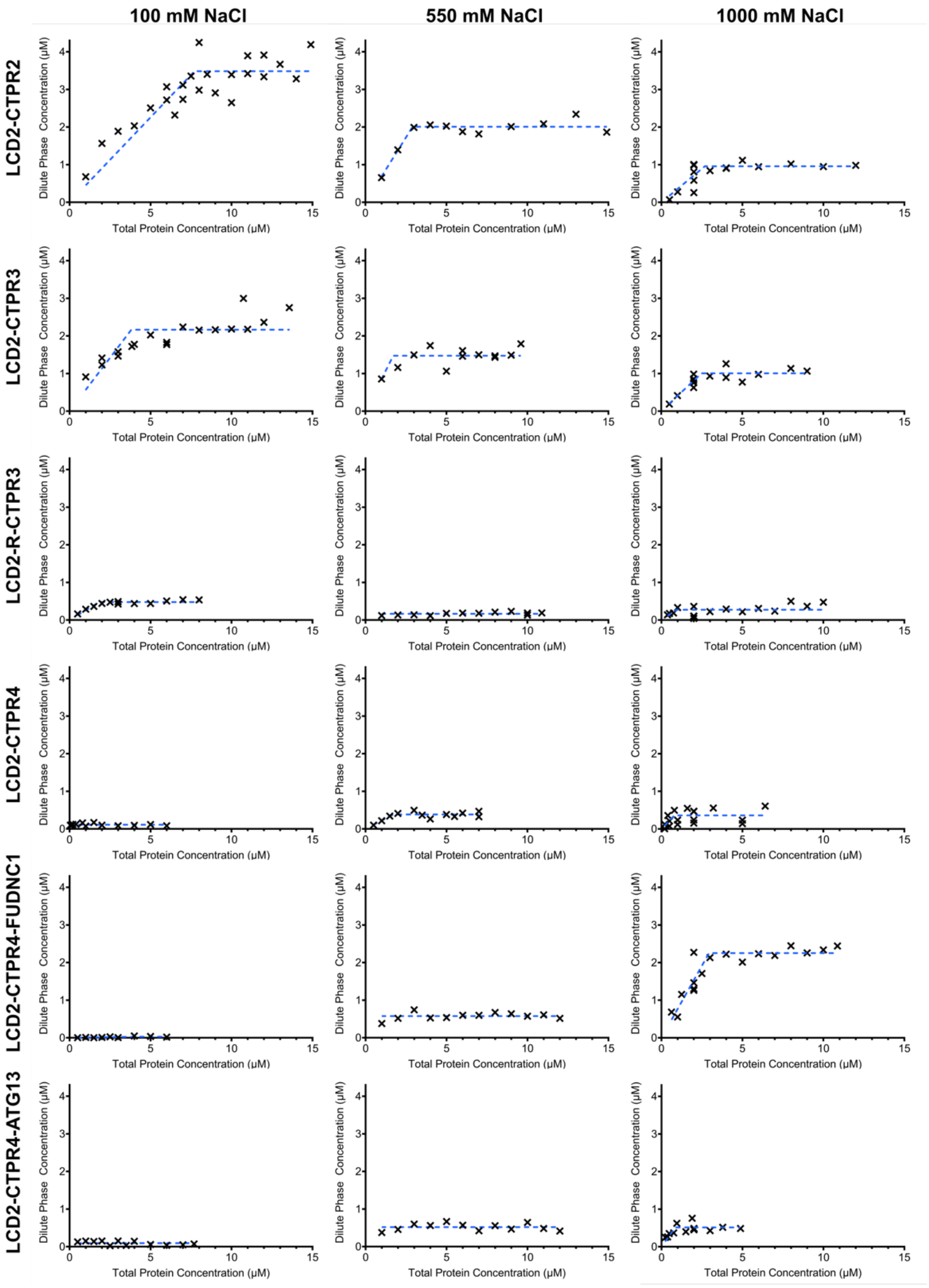


**Fig. S9: $C_{sat}$ assay for LCD2-CTPR variants.** Plots show dilute phase concentrations reach plateau at $C_{sat}$ concentration as total protein concentration increases. Proteins were in 50 mM Tris, pH 7, 300 mM urea, variable NaCl conditions.

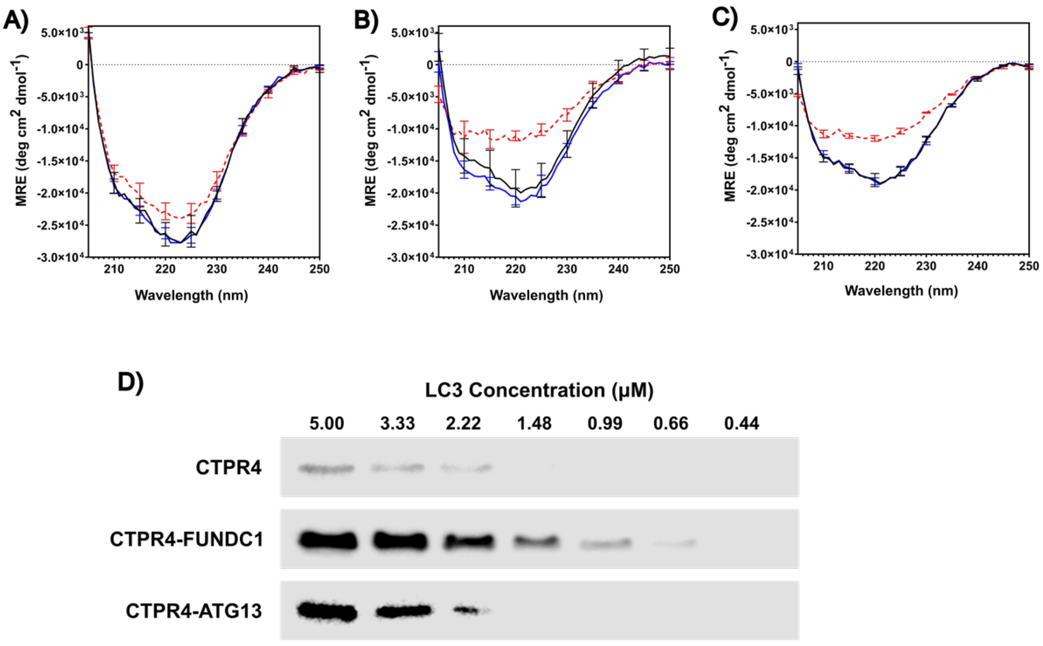


**Fig. S10: A-C) CTPR-LIR Circular Dichroism characterisation.** A) CTPR4, B) CTPR4-FUNDC1, C) CTPR4-ATG13. 5 μm of protein in 50 mM sodium phosphate, 150 mM NaCl, 0.5 mM TCEP, pH 7.5, spectra recorded from 205 to 250 nm in 0.5 nm intervals. Measured at 20°C (black), 90°C (red) and return to 20°C (blue). **D) CTPR-LIR – LC3 binding pull down.** Western blot showing the difference in capabilities of CTPR-LIR variants to pull down purified LC3 from solution at a range of concentrations.

**Table S1.** Amino acid sequences of LLPS-CTPR variants. The CTPR variant is indicated in bold and where substitutions to arginine are made, these are in blue italics. LCD1 N- and C-terminal sequences are in light blue, LCD2 N- and C-terminal sequences are in purple. Where applicable, the solvating helix is underlined. The tetracysteine-tag is indicated in green, the FUNDC1 sequence in red and the ATG13 sequence in blue. For optimised purification of full-length protein, LCD2-variants have an N-terminal His-tag, whereas LCD1-variants have a C-terminal His-tag.

| Protein name | Amino acid sequence |
|---|---|
| LCD1-CTPR3 | MGSINNNFNTNNNSNTDLDRDWKTALNIPKKDTRPQTDDVLNT<br>KGNTGS**AEAWYNLGNAYYKQGDYQKAIEYYQKALELDPNNC**<br>**CPGCCDPNNAEAWYNLGNAYYKQGDYQKAIEYYQKALELDP**<br>**RSAEAWYNLGNAYYKQGDYQKAIEYYQKALELDPRSAEAKQ**<br>**NLGNAKQKQG**KLVPVPFPIEQQSYHQQAIPQQQLPSQQQFAIP<br>PQQHHPQFMVPPSHQQQQAYPPPQMPSQQGYPPQQEHFMA<br>MPPGQSQPQYLEHHHHHH |
| LCD2-CTPR2 | MRSGHHHHHHGLVPRGSADLPQKVSNLSINNKENGGGGGKSS<br>YVPPHLRSRGKPSFERRSPKQKDKVTGGDFFRRAGRQTGNNG<br>GFFGFSKERNGGTSANYNRRGSSNYKSSGNRWVNGKHIPGPK<br>NAKLQKAELFGVHDDPDYHSSGIKFDNYDNIPVDASGKDVPEPIL<br>GS **AEAWYNLGNAYYKQGDYQKAIEYYQKALELDPNNCCPGCC**<br>**DPNNAEAWYNLGNAYYKQGDYQKAIEYYQKALELDPRSAEAK**<br>**QNLGNAKQKQG**KLGGRTRGGGGFFNSRNNGSRDYRKHGGNG |

| | |
|---|---|
| | SFGSTRPRNTGTSNWGSIGGGFRNDNEKNGYGNSNASWW |
| LCD2-CTPR3 | MRSGHHHHHHGLVPRGSADLPQKVSNLSINNKENGGGGGKSS<br>YVPPHLRSRGKPSFERRSPKQKDKVTGGDFFRRAGRQTGNNG<br>GFFGFSKERNGGTSANYNRRGSSNYKSSGNRWVNGKHIPGPK<br>NAKLQKAELFGVHDDPDYHSSGIKFDNYDNIPVDASGKDVPEPIL<br>GS AEAWYNLGNAYYKQGDYQKAIEYYQKALELDPNNCCPGCC<br>DPNNAEAWYNLGNAYYKQGDYQKAIEYYQKALELDPRSAEAW<br>YNLGNAYYKQGDYQKAIEYYQKALELDPRSAEAKQNLGNAKQ<br>KQGKLGGRTRGGGGFFNSRNNGSRDYRKHGGNGSFGSTRPR<br>NTGTSNWGSIGGGFRNDNEKNGYGNSNASWW |
| LCD2-R-CTPR3 | MRSGHHHHHHGLVPRGSADLPQKVSNLSINNKENGGGGGKSS<br>YVPPHLRSRGKPSFERRSPKQKDKVTGGDFFRRAGRQTGNNG<br>GFFGFSKERNGGTSANYNRRGSSNYKSSGNRWVNGKHIPGPK<br>NAKLQKAELFGVHDDPDYHSSGIKFDNYDNIPVDASGKDVPEPIL<br>GS AEAWYNLGNAYY*R*QGDYQ*R*AIEYYQ*R*ALELDPNNCCPGCC<br>DPNNAEAWYNLGNAYY*R*QGDYQ*R*AIEYYQ*R*ALELDPRSAEAW<br>YNLGNAYY*R*QGDYQ*R*AIEYYQ*R*ALELDPRSAEAKQNLGNAKQ<br>KQGKLGGRTRGGGGFFNSRNNGSRDYRKHGGNGSFGSTRPR<br>NTGTSNWGSIGGGFRNDNEKNGYGNSNASWW |
| LCD2-CTPR4 | MRSGHHHHHHGLVPRGSADLPQKVSNLSINNKENGGGGGKSS<br>YVPPHLRSRGKPSFERRSPKQKDKVTGGDFFRRAGRQTGNNG<br>GFFGFSKERNGGTSANYNRRGSSNYKSSGNRWVNGKHIPGPK<br>NAKLQKAELFGVHDDPDYHSSGIKFDNYDNIPVDASGKDVPEPIL<br>GSAEAWYNLGNAYYKQGDYQKAIEYYQKALELDPNNCCPGCC<br>DPNNAEAWYNLGNAYYKQGDYQKAIEYYQKALELDPNNAEAW<br>YNLGNAYYKQGDYQKAIEYYQKALELDPNNAEAWYNLGNAYYK<br>QGDYQKAIEYYQKALELDPNNKLGGRTRGGGGFFNSRNNGSRD<br>YRKHGGNGSFGSTRPRNTGTSNWGSIGGGFRNDNEKNGYGNS<br>NASWW |
| LCD2-CTPR4-ATG13 | MRSGHHHHHHGLVPRGSADLPQKVSNLSINNKENGGGGGKSS<br>YVPPHLRSRGKPSFERRSPKQKDKVTGGDFFRRAGRQTGNNG<br>GFFGFSKERNGGTSANYNRRGSSNYKSSGNRWVNGKHIPGPK<br>NAKLQKAELFGVHDDPDYHSSGIKFDNYDNIPVDASGKDVPEPIL<br>GSAEAWYNLGNAYYKQGDYQKAIEYYQKALELDPNNCCPGCC<br>DPNNAEAWYNLGNAYYKQGDYQKAIEYYQKALELDPNNAEAW<br>YNLGNAYYKQGDYQKAIEYYQKALELDPNNGGSSGNTHDDFVM<br>IDFKPADPNNAEAWYNLGNAYYKQGDYQKAIEYYQKALELDPNN<br>KLGGRTRGGGGFFNSRNNGSRDYRKHGGNGSFGSTRPRNTGTS<br>NWGSIGGGFRNDNEKNGYGNSNASWW |
| LCD2-CTPR4-FundC1 | MRSGHHHHHHGLVPRGSADLPQKVSNLSINNKENGGGGGKSS<br>YVPPHLRSRGKPSFERRSPKQKDKVTGGDFFRRAGRQTGNNG<br>GFFGFSKERNGGTSANYNRRGSSNYKSSGNRWVNGKHIPGPK<br>NAKLQKAELFGVHDDPDYHSSGIKFDNYDNIPVDASGKDVPEPIL<br>GSAEAWYNLGNAYYKQGDYQKAIEYYQKALELDPNNCCPGCC<br>DPNNAEAWYNLGNAYYKQGDYQKAIEYYQKALELDPNNAEAW<br>YNLGNAYYKQGDYQKAIEYYQKALELDPNNDYESDDDSYEVLD<br>LTEYDPNNAEAWYNLGNAYYKQGDYQKAIEYYQKALELDPNN<br>KLGGRTRGGGGFFNSRNNGSRDYRKHGGNGSFGSTRPRNTGT<br>SNWGSIGGGFRNDNEKNGYGNSNASWW |

**Table S2. Mass spectrometry results for purified LCD2-CTPR variants.**

| Protein | Expected Mass /Da | Observed Mass /Da |
|---|---|---|
| LCD2-CTPR2 | 36883.2 | 36886.5 ± 0.7 |
| LCD2-CTPR3 | 40925.6 | 40928.3 ± 2.0 |
| LCD2-RCTPR3 | 41177.7 | 41181.4 ± 6.3 |
| LCD2-CTPR4 | 43356.1 | 43360.0 ± 5.3 |
| LCD2-CTPR4-ATG13 | 45773.7 | 45777.0 ± 7.3 |
| LCD2-CTPR4-FUNDC1 | 45849.6 | 45852.2 ± 1.7 |

**Table S3.** Parameters obtained from two-state fit of the GdnHCl-induced chemical denaturation experiments for CTPR variants lacking the molecular adhesives. All variants have a tetracysteine motif between repeats 1 and 2 unless stated otherwise. CTPR2, CTPR3, R-CTPR3 and CTPR3 (no TC-tag) all have a C-terminal solvating helix.

| Protein | $D_{50\%}$ (M) | *m*-value (kcal mol$^{-1}$ M$^{-1}$) |
|---|---|---|
| CTPR2 | 2.6 ± 0.2 | 1.6 ± 0.4 |
| CTPR3 | 3.0 ± 0.3 | 1.7 ± 0.6 |
| R-CTPR3 | 3.1 ± 0.2 | 1.8 ± 0.9 |
| CTPR4 | 4.3 ± 0.2 | 2.8 ± 0.9 |
| CTPR4-ATG13 | 3.3 ± 0.3 | 1.5 ± 1.1 |
| CTPR4-FUNDC1 | 3.6 ± 0.1 | 2.1 ± 0.5 |
| CTPR3 (no TC-tag) | 3.6 ± 0.03 | 2.7 ± 0.5 |
| CTPR4 (no TC-tag) | 4.8 ± 0.1 | 3.8 ± 0.6 |

**Uncropped gels and western blots**

Supplementary Figure 1:

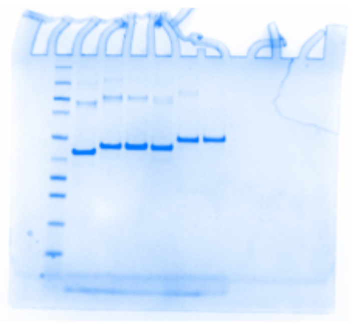

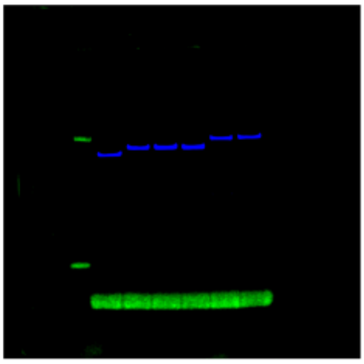

Supplementary Figure 9:

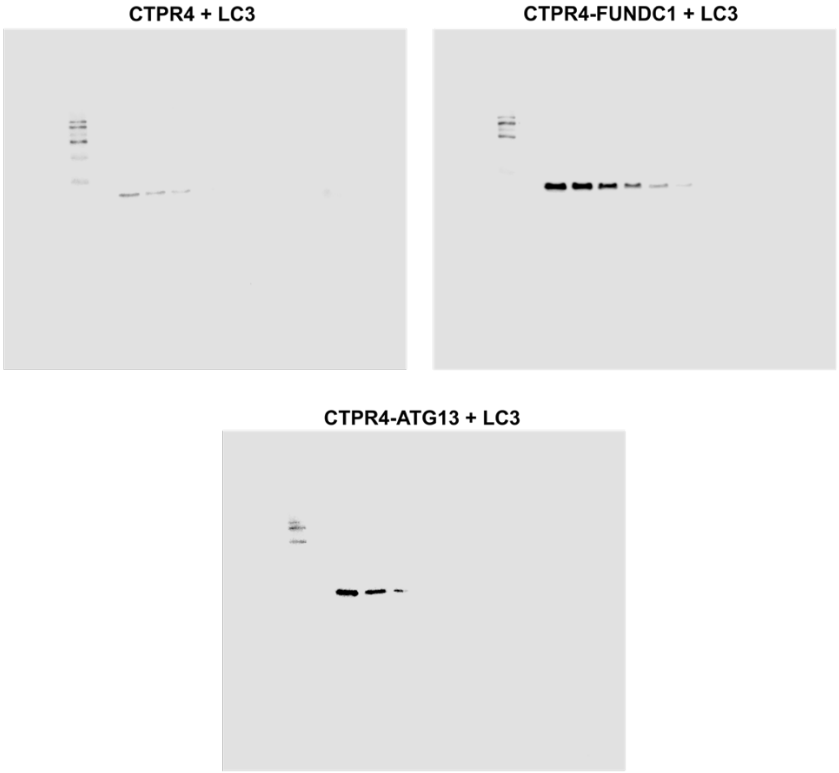
